# Integrin-like adhesin CglD confers traction and stabilizes bacterial focal adhesions involved in myxobacterial gliding motility

**DOI:** 10.1101/2023.10.19.562135

**Authors:** Nicolas Y. Jolivet, Endao Han, Akeisha M. Belgrave, Fares Saïdi, Newsha Koushki, David J. Lemon, Laura M. Faure, Betty Fleuchot, Utkarsha Mahanta, Heng Jiang, Gaurav Sharma, Jean-Bernard Fiche, Benjamin P. Bratton, Mamoudou Diallo, Beiyan Nan, David R. Zusman, Guillaume Sudre, Anthony Garza, Marcelo Nollmann, Allen J. Ehrlicher, Olivier Théodoly, Joshua W. Shaevitz, Tâm Mignot, Salim T. Islam

## Abstract

Integrins are crucial for eukaryotic cell attachment and motility within the extracellular matrix (ECM) via focal-adhesion formation, with their evolutionary emergence important for the development of multicellularity. Intriguingly, single gliding cells of the predatory deltaproteobacterium *Myxococcus xanthus* form bacterial focal-adhesion (bFA) sites; therein, helically-trafficked motors become immobilized at anchored locations through Glt apparatus association with cell-surface integrin αI-domain-like adhesin CglB. Using traction-force, bead-force, and total internal reflection-fluorescence microscopies combined with biochemical approaches, we herein identify the von Willebrand A domain-containing cell-surface lipoprotein CglD to be a β-integrin-like outer-membrane lipoprotein that functionally associates with and anchors the trans-envelope Glt–CglB gliding apparatus, stabilizing and efficiently anchoring this assembly at bFAs. Calcium dependence governs CglD importance, consistent with its integrated ECM eukaryotic cartilage oligomeric matrix protein domains. CglD thus confers mechanosensory and mechanotransductory capabilities to the gliding apparatus, helping explain bFA-mediated trans-envelope force transduction, from inner-membrane-embedded motors to the cell surface.

## Introduction

Cellular motility on surfaces necessitates complex interactions between the cell and the underlying substratum across all biological kingdoms. In metazoans, translocating cells adhere to the extracellular matrix (ECM) via nucleation of integrin proteins linked to the internal actomyosin network^1^. Integrins are composed of an α and a β subunit, with half of all α variants, and all β variants, elaborating I-domains containing a von Willebrand A (VWA) module that binds specific ligands^2^; in turn, integrin adhesion to the ECM is mediated by their interaction with soluble ECM proteins such as cartilage oligomeric matrix protein (COMP) (**Fig. 1**). Such adherence generates eukaryotic focal-adhesion (eFA) sites; these sites do not move (relative to the substratum) and transduce motor forces via induction of local traction, thus mediating cell motility relative to fixed points^3^. In bacteria such as Gram-negative *Myxococcus xanthus*, individual cells are able to glide on surfaces (without external appendages such as flagella or pili) using motorized (Agl) substratum-coupled gliding transducer (Glt) complexes that are transported towards the lagging cell pole^4,5^; similar to the abovementioned metazoan cells, these complexes in *M. xanthus* remain stationary relative to the substratum in a gliding cell and form bacterial focal adhesion (bFA) sites^6^.

**Fig. 1.**
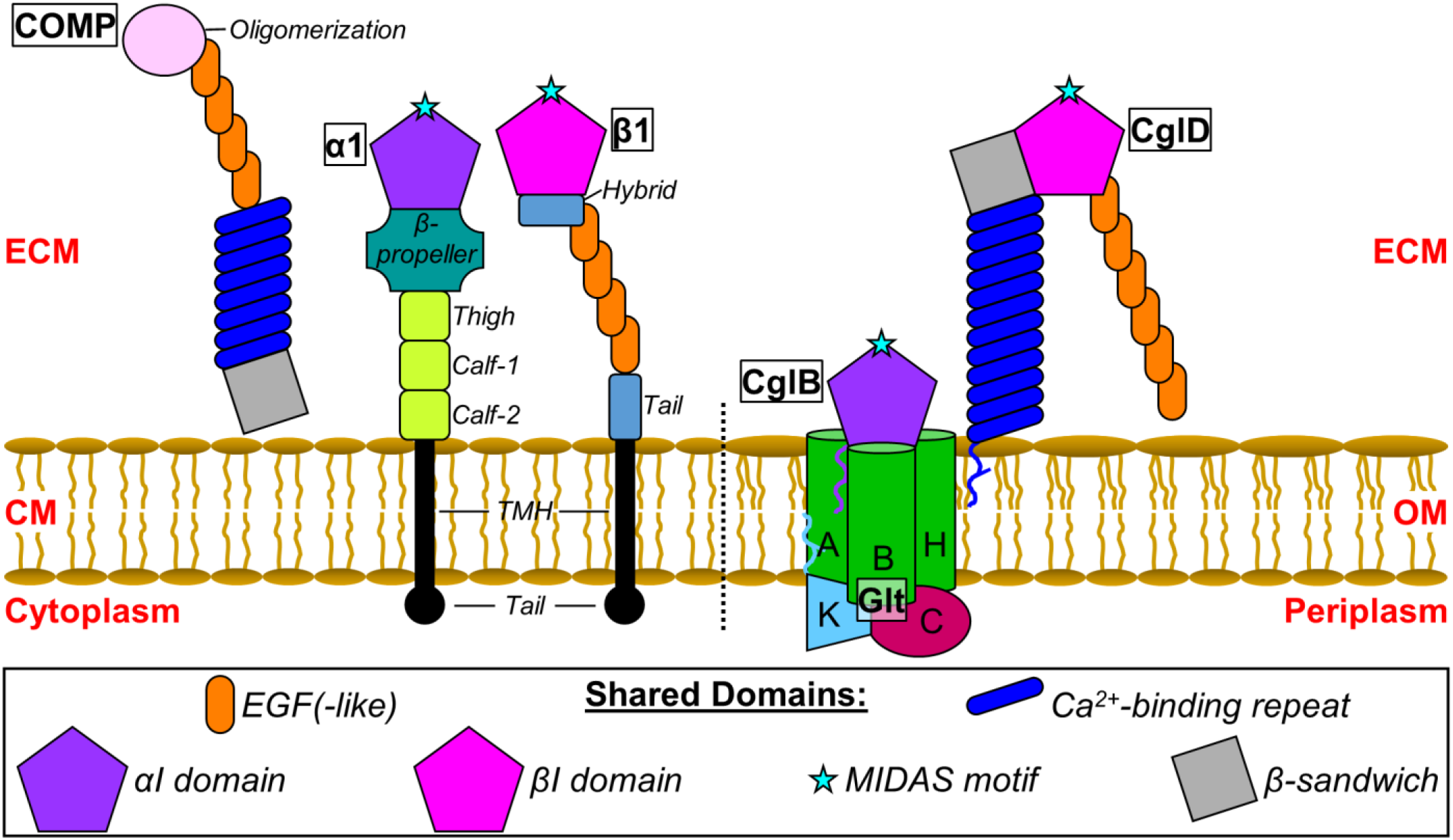
Domain schematic of eukaryotic integrin (-associated) proteins and proposed analogies with *M. xanthus* Cgl proteins. *Left:* Domains of soluble COMP (cartilage oligomeric matrix protein) as well as cytoplasmic membrane (CM)-inserted α1 and β1 integrin subunits associated with the eukaryotic extracellular matrix (ECM). *Right:* Domains analogous to those found in integrins for *M. xanthus* outer-membrane (OM) lipoproteins CglB and CglD, associated with the OM module of the Glt trans-envelope complex. GltA/B/H are integral OM β-barrels. GltK is periplasmically-oriented OM lipoprotein. GltC is an OM-associated soluble periplasmic protein. Shared domains between the various eukaryotic and prokaryotic proteins have been indicated.

*Myxococcus xanthus* is a social predatory soil deltaproteobacterium with a complex developmental cycle^7^. Under nutrient-limiting conditions, vegetative cells in a swarm biofilm aggregate, differentiate, and form multicellular spore-filled fruiting bodies. This complex life cycle is modulated by the interplay between several secreted polysaccharides and the motility of cells at the group and individual levels^8–10^. Type IV pilus extension-and-retraction is responsible for the former, while as described, the latter is mediated by Agl–Glt-dependent gliding.

In gliding *M. xanthus* cells, the motorized trans-envelope Agl–Glt complex^11,12^ is assembled at the leading pole, and is transported towards the lagging pole along a trajectory matching a right-handed helix^4^. Upon reaching the ventral side of the cell in contact with the substratum, the motorized Agl–Glt apparatus becomes coupled to the substratum via unmasking of the αI VWA domain-containing adhesin CglB, which otherwise remains loaded in the outer-membrane (OM) module of the gliding complex until engaged^5^ (**Fig. 1**). The OM module of the gliding apparatus is a hetero-oligomeric complex composed of the integral OM β-barrels GltA/B/H and periplasmically-oriented OM lipoprotein GltK, which along with OM-associated protein GltC^5,13^ form a complex that recruits and shields the surface-localized adhesin CglB ^5^. Engagement of CglB results in Agl–Glt complex immobilization at bFA sites, allowing for force transduction across the cell envelope and gliding locomotion relative to the fixed bFA^5^.

Several factors were identified >45 years ago as important for gliding when random mutagenesis screens revealed 5 classes of “conditional gliding” (*cgl*) mutations (*cglB/C/D/E/F*) that, in isolation, rendered cells gliding-null; however, mixing one class of these gliding-null mutant cells with another (e.g. *cglB* + *cglC*) resulted in a transient restoration (“stimulation”) of gliding motility across the entire population^14^. These data suggested the missing factors between mutant classes could be physically transferred between cells and integrated into the defective gliding mechanism to transiently restore single-cell locomotion. The *cglB/C/E/F* factors were later identified as specific genes encoding OM(-associated) proteins of the gliding apparatus (CglB/GltK/GltH/GltF, respectively)^11,15^. However, the role of CglD in *M. xanthus* physiology has remained contentious, with conflicting reports as to its requirement and function; though the initial randomly-generated mutant was reported to be gliding-null^14^, a subsequent clean gene-deletion mutant still resulted in gliding-capable cells, while a *cglD* missense mutation had a stronger gliding defect^15^. CglD has thus been proposed to have both an activation and an inhibition function in gliding motility^15^.

In this study, we reveal CglD to be a cell-surface β-integrin-like lipoprotein with COMP-like Ca^2+^-binding capacity. This bacterial protein with structural homologies to eukaryotic ECM-binding components at eFAs is shown to analogously confer traction to the substratum in gliding bacterial cells, impacting bFA formation, stability, and positioning. In turn, this drastically influences both single-cell and community-level events that are essential to multicellular outcomes.

## Results

### CglD is a β-integrin-like lipoprotein

To elucidate the contribution of CglD to the complex physiology of *M. xanthus*, we first set out identify structural motifs that could help explain its function. Fold-recognition analysis of CglD (1130 aa; predicted MW: 117 kDa) using HHpred revealed distinct high-confidence structural matches of the N- vs. C-terminal halves of the protein. CglD residues 40 – 430 were matched to the Ca^2+^-binding domain of human cartilage oligomeric matrix protein (COMP, i.e. thrombospondin type-5; PDB 3FBY_B)^16^ (**Fig. 2A**). COMP is a secreted glycoprotein that is engaged by integrins^17^ and impacts cellular attachment to, and structuration of, the extracellular matrix (ECM) in humans^17–23^. The identified dodeca-repeating motif DxDxDG^15^ (**Fig. 1**) would be consistent with a Ca^2+^-dependence for the protein. Calculation of the 3D structure of CglD via AlphaFold2 (**fig. S1A-C**) revealed this putative Ca^2+^-binding domain to form a predicted (largely unstructured) globular domain (**Fig. 2B**), consistent with its COMP structural homologue (**Fig. 2A**).

**Fig. 2.**
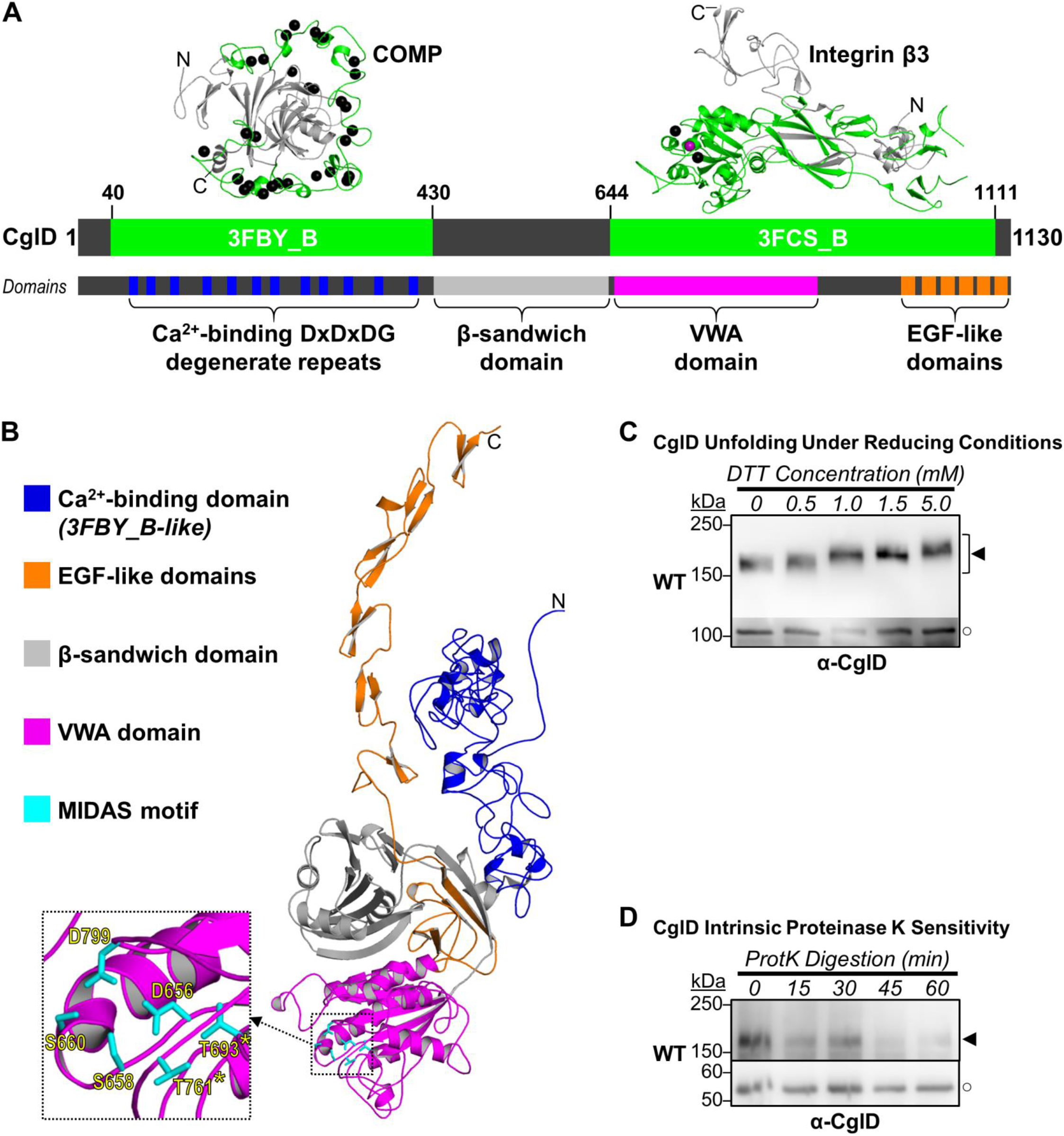
CglD contains integrin-like VWA and Ca^2+^-binding sites. **(A)** Domain organization within *M. xanthus* CglD. Regions of CglD structural homology with X-ray crystal structures of COMP (PDB: 3FBY_B) and integrin β3 (PDB: 3FCS_B) are represented in *neon green* (as determined via HHpred fold recognition). *Black spheres* depicted in the template structures are co-crystalized Ca^2+^ ions. The various domains have been coloured as follows: Ca^2+^-binding domains (*blue*); EGF-like domains (*orange*); β-sandwich (*grey*); VWA domain (*magenta*); MIDAS motif (*cyan*). **(B)** Alphafold model of CglD protein, with domains coloured as in Panel A Inset: magnified view of the MIDAS motif, with putative amino acids indicated. **(C)** α-CglD Western blot of WT whole-cell extracts treated with increasing concentrations of DTT to break disulphide bonds. The lower, darker zone on the blot corresponds to the section of the same blot image for which the contrast has been increased to highlight lower-intensity protein bands. Legend: ◄, full-length CglD; ○, loading control (labelled non-specifically by α-CglD pAb). **(D)** Intact WT cells resuspended in TPM buffer and digested with exogenous Proteinase K. Aliquots of the digestion mixture were removed at 15-min intervals and TCA-precipitated to stop digestion. The lower zone on the blot corresponds to a lower section of the same blot image for which the contrast has been increased to highlight lower-intensity protein bands. Legend: ◄, full-length CglD; ○, non-specific loading control.

Conversely, CglD amino acids 644 – 1111 were identified as a match to the βI domain of human integrin β3 (i.e. CD61; PDB 3FCS_B) (**Fig. 2A**); βI modules are ubiquitous in β integrins and include a VWA domain with a metal ion-dependent adhesion site (MIDAS) motif implicated in adhesion^2^. MIDAS motifs are a discontinuous structural element (Asp-x-Ser-x-Ser…Thr…Asp) that coordinate a divalent cation (e.g. Ca^2+^/Mn^2+^/Mg^2+^), structurally modifying VWA domains upon binding of their ligand(s) to generate a high-affinity conformation toward the ligand(s). Based on the predicted structure for CglD, D656, S658, S660, T693/761, and D799 constitute the putative MIDAS amino acids (**Fig. 2B**); while the former three residues were previously predicted^15^, the remaining residues were identified based on their spatial proximity and orientation towards the canonical DxSxS tract. A mutant strain of *M. xanthus* encoding a chromosomal D656N variant of CglD was found to be compromised for gliding motility-dependent swarm-edge flare formation^15^, consistent with the MIDAS motif being functionally important for CglD.

Intriguingly, CglD was found to be a Cys-rich protein (46 of 1130 amino acids = 4.1%), with 18 predicted intra-protein disulphide bonds stabilizing various structural loops (**fig. S1D**). Titration of whole-cell lysates with reducing agent followed by α-CglD Western immunoblot analysis revealed CglD-specific bands shifting from faster to slower-migrating protein species, consistent with disulphide-dependent conformational stability of CglD (**Fig. 2C**). Eleven of these disulphide bonds were located in repeating predicted anti-parallel domains resembling epidermal growth factor (EGF) in humans (**Fig. 2B**, **S1E**). Conversely, this unfolding of CglD was not impacted by the amount of Ca^2+^ to which the protein was exposed during cell growth (**fig. S2**), suggesting that any Ca^2+^-binding capacity of the β-integrin-like protein may not serve to stabilize its own structure.

Given its’ (i) overall domain arrangement, (ii) structural homologies to known human counterparts, and (iii) denaturation profiles, we conclude that CglD is a β-integrin-like protein with an integrated COMP module.

### CglD is exposed at the cell surface

Lipoproteins in the OM were historically thought to localize to its periplasmic leaflet; however, cell-surface exposure of various lipoproteins is becoming more widely acknowledged^24,25^, including with our recent determination of surface localization for the principal gliding adhesin lipoprotein CglB^5^. Given the calculated integrin-like structure of CglD (**Fig. 2B**), we hypothesized that it too is exposed at the cell surface. We initially attempted immunolabelling of CglD (with α-CglD pAb) on live cells, but various fluorescent clusters were detected on both WT and Δ*cglD* cells. To overcome this ambiguity, we adopted an analogous approach to that used to test CglB surface exposure in the presence/absence of OM-module Glt proteins^5^. I.e. intact WT cells were digested with Proteinase K over the course of 60 min; aliquots removed at regular intervals revealed a steady decline in full-length CglD signal (with no visible accumulation of pAb-reactive degradation products) (**Fig. 2D**). These data are consistent with β-integrin-like CglD being a surface-exposed lipoprotein.

### CglD modulates *M. xanthus* community structuration/behaviour

As integrins in humans serve to interact with cells/substrata and structure the ECM, we set out to probe comparable community-level outputs in *M. xanthus*. Swarms of *M. xanthus* spreading on “soft” (0.5% w/v) agar plates form stratified biofilms of cells surrounded by ECM polysaccharides^9^. In the absence of β-integrin-like CglD, swarm expansion was consistently found to be negatively impacted; this community-level effect was not due to a potential defect in gliding motility at the single-cell level as swarm expansion in a gliding-deficient Δ*cglB* strain (compromised for surface-coupling of the gliding machinery) was instead found to be augmented relative to WT (**Fig. 3A,B**), consistent with a previous report^26^. Similarly, developmental progression on minimal media requiring cell–cell aggregation was more drastically affected in the absence of CglD compared to the absence of CglB, further supporting a role for CglD in forming inter-cell connections in dense populations.

**Fig. 3.**
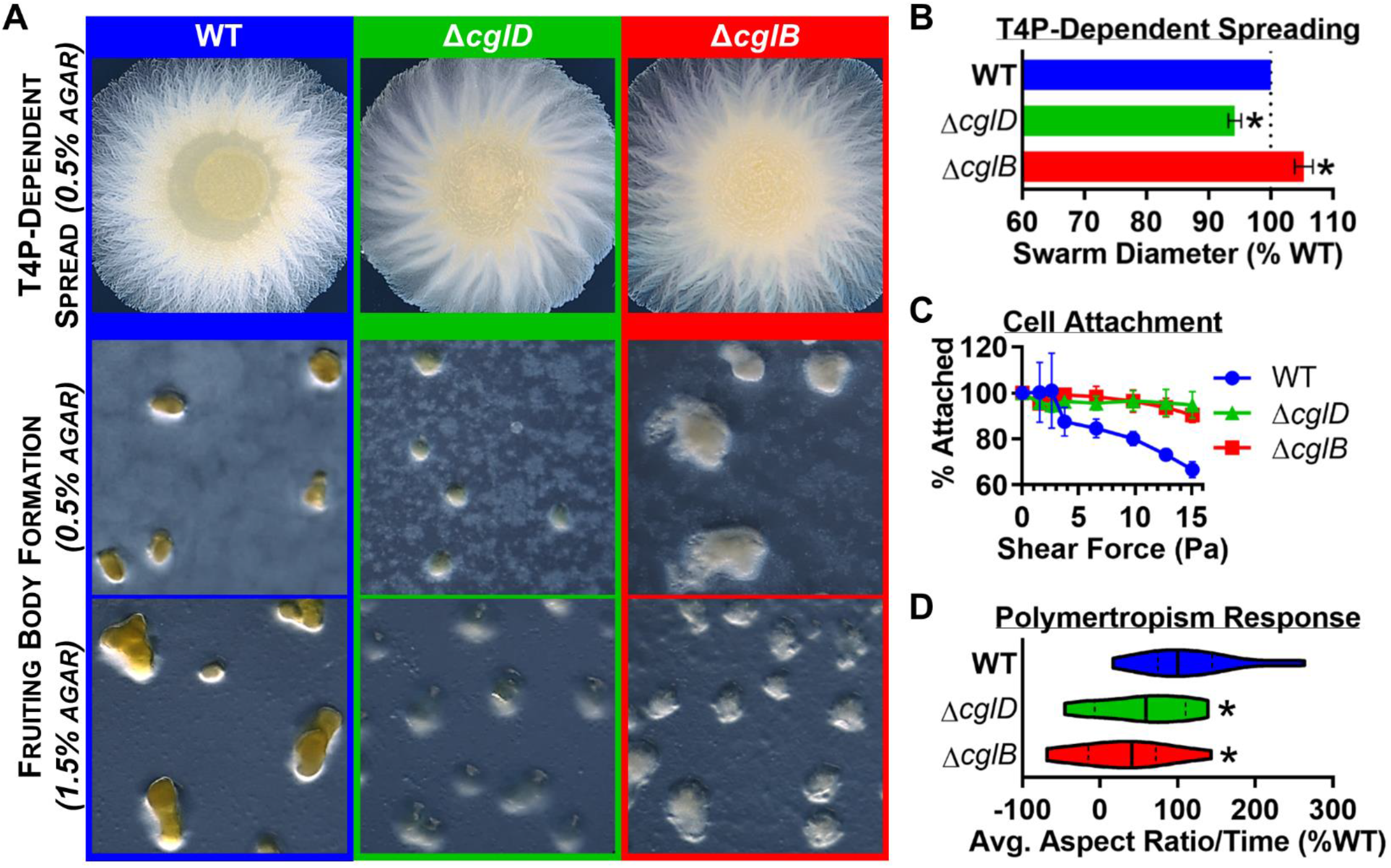
Impact of CglD deficiency on multicellular behaviours. **(A)** Macroscopic phenotype comparison between WT, Δ*cglD* and Δ*cglB* strains. *Top row:* T4P-dependent swarm spreading on CYE 0.5% agar plates. *Middle and bottom rows*: Fruiting body formation on CF 0.5% and 1.5% agar plates (respectively) after 72 h at 32 °C. **(B)** Swarm diameter measurements for T4P-dependent spreading, normalized to WT (n = 5). Both Δ*cglD* and Δ*cglB* swarms displayed significantly different (*) swarm diameters compared to WT, as determined via Wilcoxon signed-rank test set relative to a reference value of 100 (*p* < 0.05). **(C)** Swarm integrity analysis as determined via rheometric shearing of adhered fluorescent cell strains. Fluorescence values at all shear forces were normalized to the intensity from the initial image acquired prior to shear-force application. Each shear force point indicated is the mean of 3 biological replicates (± SEM). Increasing shear forces were applied for a duration of 1 min after which the force was increased to the next level via faster rotation of the rheometer arm. **(D)** Polymertropism response determined by calculating the average aspect ratio of a swarm over time, normalized to the WT control strain run at the same time. on time and normalizing it on the WT. Both Δ*cglD* and Δ*cglB* displayed significantly different dataset distributions (*) compared to the values for WT, as determined via Mann-Whitney test (*p* < 0.05). Indicate figure parts with bold capital letters: (**A**), (**B**), etc.

Swarm cohesiveness was also impacted by CglD, with larger clumps of fluorescent cells in WT swarms being sloughed off in response to increasing shear force applied by a rheometer (**Fig. 3C**, **S3**); however, as the same swarm cohesiveness phenotype was shown to depend on the principal gliding adhesin CglB (**Fig. 3C**), this would suggest a role for CglD in gliding motility, consistent with previous reports^14,15^, with gliding motility potentially rendering swarms more dynamic and hence less stable. Similarly, community-level responses to mechanical substratum changes were affected in the absence of CglD, as assayed via polymertropism response (i.e. changes in swarm aspect ratio) (**Fig. 3D**). Polymertropism is a gliding- and glycocalyx-dependent phenomenon^9,27^ in which swarms preferentially spread in an east–west direction, in response to aligned substratum polymers caused by north–south compression of the underlying agar matrix^27–29^. While the absence of CglB resulted in a severely-compromised polymertropism response, swarms lacking CglD displayed an intermediate polymertropism phenotype (**Fig. 3D**), again implicating CglD in gliding motility (albeit in a non-essential capacity).

Taken together, the above-described data are consistent with β-integrin-like CglD playing roles in both community-level inter-cell interactions as well as gliding motility outcomes.

### CglD promotes (but is not required for) gliding motility on deformable substrata

As the *cglD* locus was originally identified through its conditional importance for gliding motility, we next set out to reconcile the disparate datasets in which CglD has been postulated to either be required^14^ or not^15^ for gliding motility. We first compared gliding-dependent swarm-edge flare formation for WT and Δ*cglD* cells across diverse matrices, each characterized by a distinct elastic modulus (i.e. resistance to deformation). For agar (1.5% w/v), carrageenan (1.5% w/v), and gellan (0.6% w/v) matrices, gliding flares were clearly detected for both strains already after 7 h (**Fig. 4A**). Incidentally, flares on gellan were considerably more prominent/noticeable compared to those on either carrageenan or (canonical) agar substrata (**Fig. 4A**), suggesting that gellan-based matrices may be a superior alternative for the examination of myxobacterial gliding-flare formation. For both strains, individual and groups of cells were found to follow previously-carved troughs in the deformable matrices, further supporting the notion of sematectonic stigmergy^9,30^ — i.e. behavioural coordination within a population without direct interactions between individuals, accomplished via physical modifications of the local environment — being responsible for the eponymous “trail following” phenotype of myxobacteria atop agar matrices^31,32^ (**Fig. 4A**). At the level of individual cells, those lacking CglD were found to glide on the various substrata at slower speeds (**Fig. 4B**) with less-frequent reversals of gliding direction (**Fig. 4C**).

**Fig. 4.**
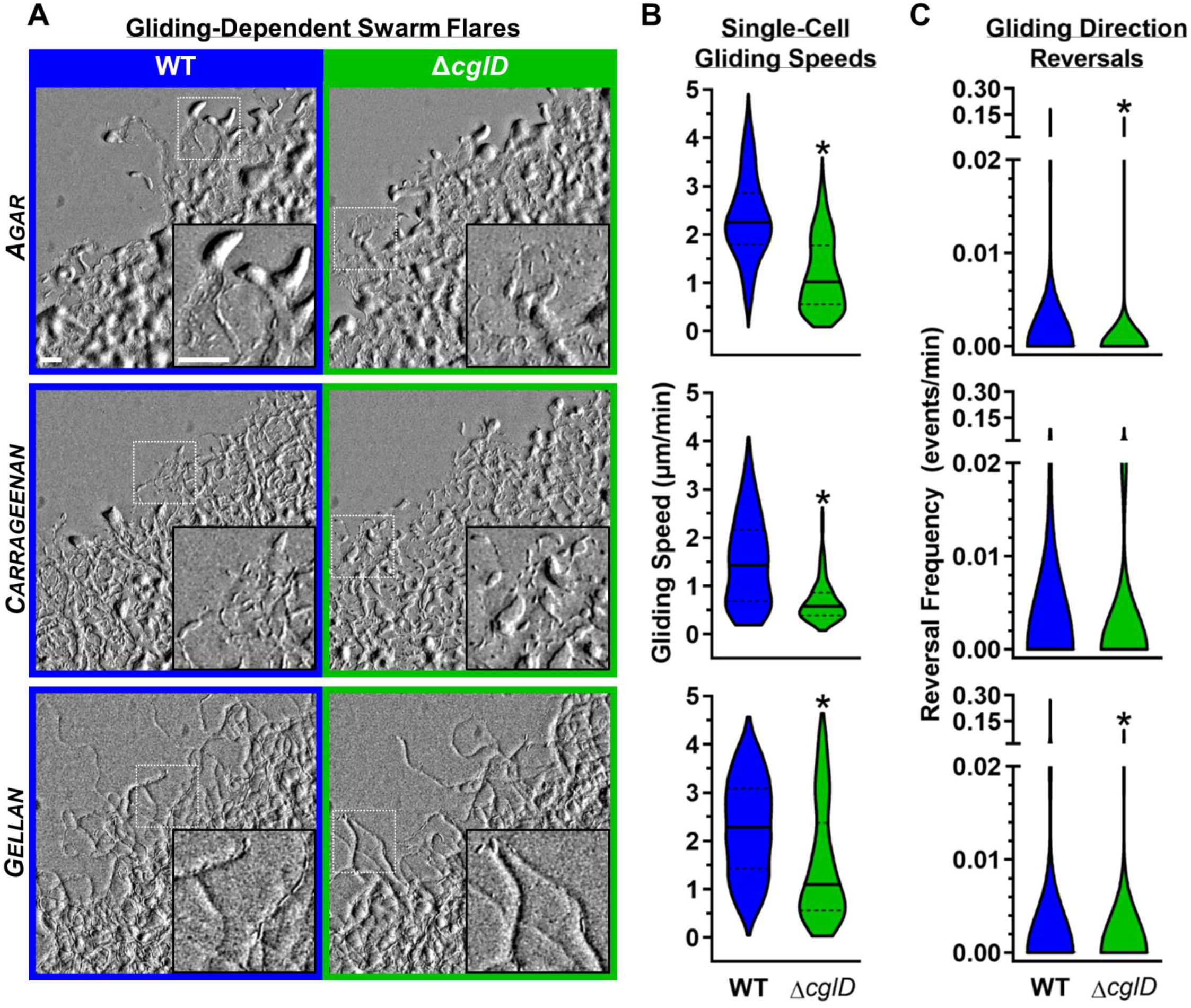
CglD deficiency impacts gliding motility across multiple substrata. Each row referred to a specific gelling agents used: Agar 1.5 %, Carrageenan 1.5% and gellan 0.6% respectively from top to bottom. **(A)** Gliding-dependent swarm-edge flares on CYE substrata solidified with 1.5% agar, 1.5% carrageenan, or 0.6% gellan. The insets represent magnified views of the indicated areas (*white dashed boxes*) showing deep furrows carved out by flare-leading cells, which can be followed by subsequent cells via sematectonic stigmergy. Scale bars at both magnifications: 50 µm. **(B)** Violin plots of single-cell gliding-*event* speeds for WT and Δ*cglD* cells on pads solidified with different gelling agents across 3 biological replicates (agar: n_WT_ = 1534 events, n_Δ*cglD*_ = 1237 events; carrageenan: n_WT_ = 233 events, n_Δ*cglD*_ = 272 events; gellan: n_WT_ = 1826 events, n_Δ*cglD*_ = 627 events). The lower and upper boundaries of the plots correspond to the minimum and maximum values of the dataset, with the 25^th^ and 75^th^ percentiles displayed (*dashed black lines*). The median (*solid black line*) of each dataset is indicated. Asterisks denote datasets displaying statistically significant (*) differences in distributions (*p* < 0.0001) between WT and Δ*cglD* cells, as determined via unpaired two-tailed Mann– Whitney tests. **(C)** Violin plots of reversal events per minute for tracked WT and Δ*cglD* cells (see Panel B) on pads solidified with agar, carrageenan, or gellan across 3 biological replicates (agar: n_WT_ = 1534 events, n_Δ*cglD*_ = 1237 events; carrageenan: n_WT_ = 233 events, n_Δ*cglD*_ = 272 events; gellan: n_WT_ = 1826 events, n_Δ*cglD*_ = 627 events). The lower and upper boundaries of the plots correspond to the minimum and maximum values of the dataset, with the 25^th^ and 75^th^ percentiles as well as the median not distinguishable due to skewing of the data by non-reversing cells. Asterisks denote datasets displaying statistically significant (*) differences in distributions (*p* < 0.0001) between WT and Δ*cglD* cells, as determined via unpaired two-tailed Mann–Whitney tests.

The only instance in which we found Δ*cglD* cells unable to manifest gliding-dependent flares compared to flare-forming WT cells was related to the drying conditions used for the agar matrix. Namely, freshly-poured “hard” (1.5% w/v) agar plates left to cool uncovered in the biohood for increasing 10-min intervals prior to covered drying on benchtop (2 h) and spotting of cells, sealing of the plate, and incubation for 24 h. These parameters were able to support gliding-flare formation in WT cells, but not in Δ*cglD* cells. Conversely, identical plates allowed to dry uncovered for longer in the biohood were indeed able to support gliding-flare formation in both strains (**fig. S4**). As such, the previously reported^14^ absolute requirement for CglD in agar-based gliding may have been (at least partially) due to the hydration state of the agar matrix being used.

### CglD is required for Ca^2+^-dependent gliding motility on rigid substrata

With the high propensity of its β-integrin-like tertiary structure to bind calcium ions (**Fig. 2A,B**), we next examined the contribution of Ca^2+^ to CglD-modulated gliding motility. To precisely control the amount of calcium present, Ca^2+^-dependent single-cell motility was established in polydimethylsiloxane (PDMS) microfluidic chambers via use of non-deformable chitosan-functionalized glass substrata; as the concentration of Ca^2+^ in the chitosan preparation buffer increased, so too did the gliding speed of cells (**Fig. 5A**). Cells lacking CglD were found to be severely compromised for Ca^2+^-dependent gliding on chitosan-functionalized glass (**Fig. 5B**), demonstrating the essentiality of the β-integrin-like surface protein in this context. Thus, while CglD is dispensable for single-cell gliding on deformable matrices, the β-integrin-like protein is necessary for single-cell locomotion on a rigid substratum in a calcium-dependent manner.

**Fig. 5.**
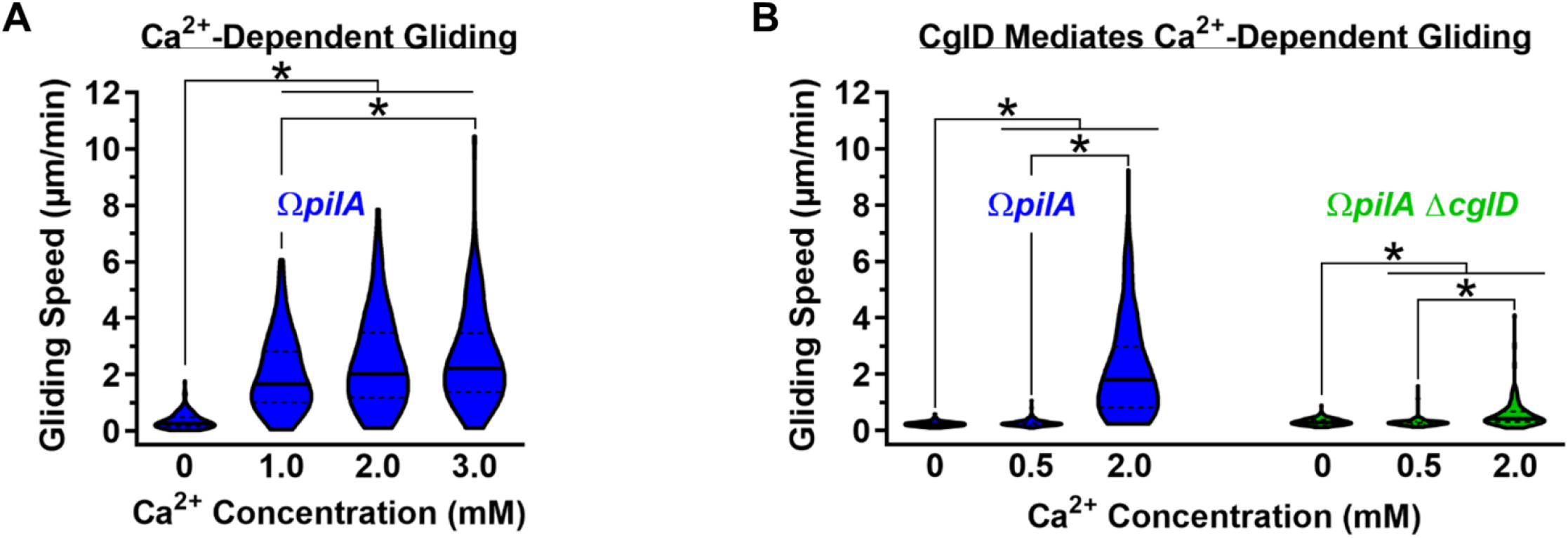
Single-cell gliding motility on non-deformable chitosan-functionalized glass is Ca^2+^- and CglD-dependent. **(A)** Violin plots of single-cell gliding event speeds for Ω*pilA* cells in PDMS microfluidic chambers atop chitosan-functionalized glass slides. CaCl_2_ was present in increasing concentrations in separate microfluidic channels. **(B)** Violin plots of single-cell gliding event speeds for Ω*pilA* and Ω*pilA* Δ*cglD* cells on glass coverslips coated with chitosan in the presence of increasing concentrations of CaCl_2_ (n = 115 cells). For Panels A and B, the lower and upper boundaries of the plots correspond to the minimum and maximum values of the dataset, with the 25th and 75th percentiles displayed (*dashed lines*). The median (*solid black line*) of each dataset is indicated. Asterisks denote datasets displaying statistically significant differences in distributions (*p* < 0.0001) between various concentrations (*) as determined via unpaired two-tailed Mann–Whitney tests.

### CglD confers traction to gliding cells

As eukaryotic integrins are widely known to exert traction forces on the substratum under a translocating cell^33^, we postulated that β-integrin-like CglD may contribute to single-cell *M. xanthus* gliding in a comparable manner. To test this hypothesis, we undertook traction force microscopy (TFM) studies^34^ optimized for single gliding cells of various strains. TFM depends on traction-induced displacement of fluorescent particles in a temporarily deformable matrix below a translocating cell; as such, an elastic substratum capable of “springing back” following cell passage and deformation (unlike agar, carrageenan, or gellan [**Fig. 4A**]) is required. For this reason, we employed an elastic polyacrylamide (PAA) matrix coated in Ca^2+^-infused chitosan. This substratum was indeed capable of supporting gliding motility in single cells encoding the full complement of gliding machinery genes, whereas cells lacking CglB (i.e. the principal adhesin of the gliding system) were expectedly gliding deficient. In contrast, cells lacking CglD were still able to glide on this substratum, albeit at a severely-reduced capacity (**Fig. 6A**).

**Fig. 6.**
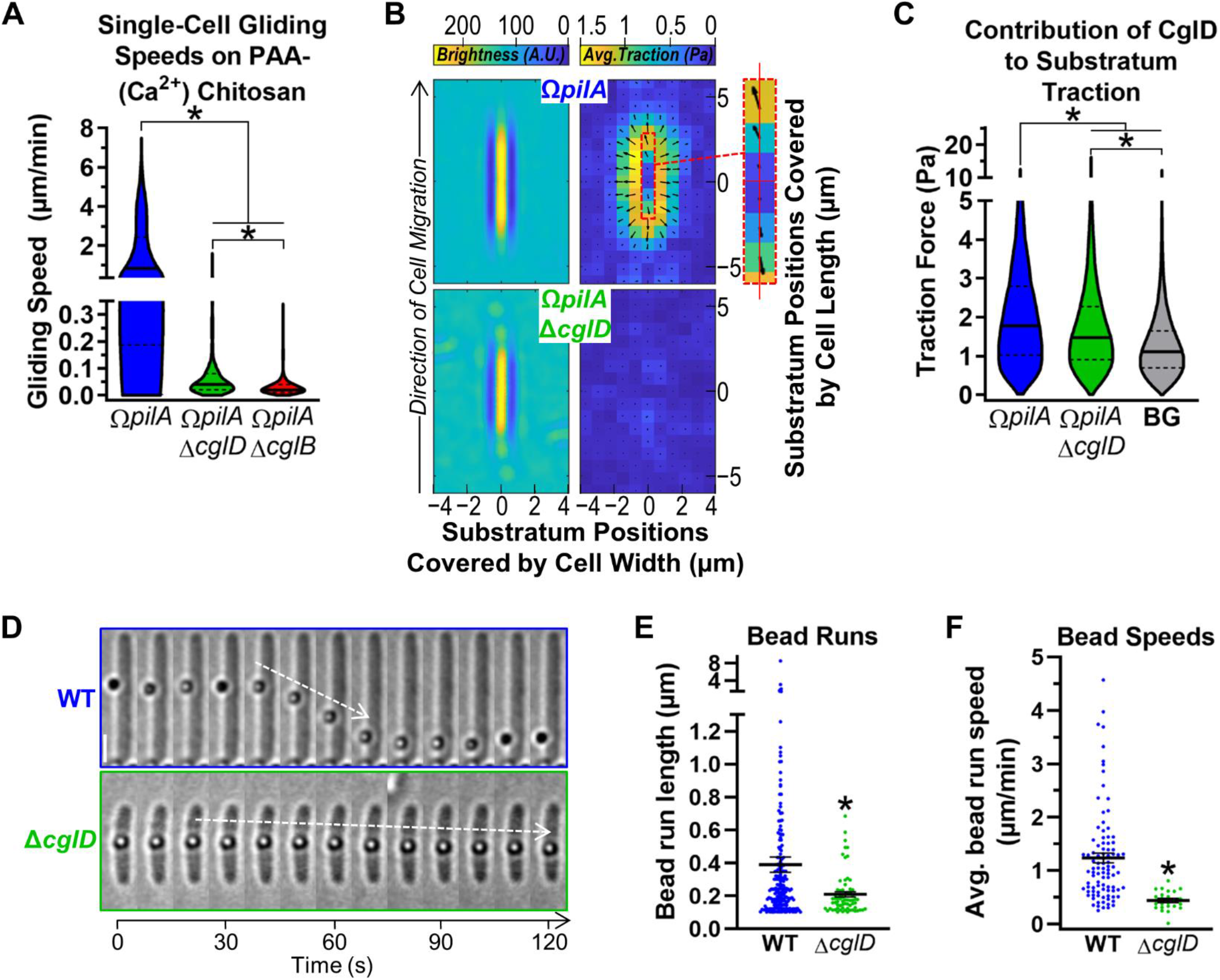
CglD engages the gliding substratum and transported cell-surface cargo. **(A)** Violin plots of single-cell gliding event speeds for Ω*pilA*, Ω*pilA* Δ*cglD*, and Ω*pilA* Δ*cglB* cells on a polyacrylamide (PAA) substratum coated with chitosan in the presence of CaCl_2_. The lower and upper boundaries of the plots correspond to the minimum and maximum values of the dataset, with the 25th and 75th percentiles displayed (*dashed black lines*). The median (*solid black line*) of each dataset is indicated. Asterisks denote datasets displaying statistically significant differences (*) in distributions (*p* < 0.0001) between each mutant strain and WT, as determined via unpaired two-tailed Mann–Whitney tests. **(B)** Traction force applied on the substratum under a gliding *M. xanthus* cell as determined via traction force microscopy. *Left-side panel:* Heat map of average cell positioning during tracked motility run, displayed as average brightness for a particular position (measured in arbitrary units, A.U.). *Right-side panel:* Heat map of the average traction force magnitude applied under a gliding cell. Arrows originating from various squares indicate the average direction of the applied traction force by the gliding cell. *Dashed red box:* Magnified view of the traction force readout immediately under CglD-containing gliding cells, indicating a horizontal skew to the direction of the applied forces. **(C)** Violin-point representing the traction forces exerted on the substratum by gliding cells with(out) CglD. The background (BG) signal is displayed to denote the baseline threshold for the TFM readings. The lower and upper boundaries of the plots correspond to the minimum and maximum values of the dataset, with the 25th and 75th percentiles displayed (*dashed black lines*). The median (*solid black line*) of each dataset is indicated. Asterisks denote datasets displaying statistically significant differences (*) in distributions (*p* < 0.0001) between each strain as determined via unpaired two-tailed Mann–Whitney tests. **(D)** Montage of the trafficking phenotypes of surface-deposited polystyrene beads on WT vs. Δ*cglD* cells. **(E)** Lengths of trafficked polystyrene bead runs (> 0.1 µm) along immobilized live *M. xanthus* cells. Images from 10-s intervals were analyzed. The distributions of the two datasets are significantly different (*), as calculated via unpaired two-tailed Mann-Whitney U-test (*p* = 0.0010). **(F)** Average speeds of trafficked polystyrene beads during bead runs. The distributions of the two datasets are significantly different (*), as calculated via unpaired two-tailed Mann-Whitney test (*p* < 0.0001).

Analysis of the distribution of traction force under a gliding cell relative to its surroundings indicated that more traction was exerted at the leading pole than at the lagging pole in cells expressing CglD, consistent with the formation of bFA sites at the leading pole; in contrast, cells deficient for CglD were only able to exert minimal traction on the substratum above background levels (**Fig. 6B,C**), providing a rationale for the considerably slower gliding speed in the latter cells (**Fig. 6A**). As force is a vector quantity, in addition to its magnitude, we also examined the directionality of the applied traction force. Importantly, immediately under the leading halves of advancing cells, traction forces were exerted with a leftward orientation, while under the lagging halves of advancing cells, traction forces were applied with a rightward bias (**Fig. 6B, *inset***).

Taken together, these data indicate that CglD contributes traction forces to the substratum under gliding cells. Moreover, the directionality of these forces provides independent corroboration that gliding *M. xanthus* cells undergo helical rotation of the cell along its long axis^4^ while gliding.

### CglD is directly involved in the gliding mechanism

The TFM data for strains with(out) CglD could be explained by one of two scenarios:

i. CglD serves as a general cell-surface adhesin that non-specifically helps bring the cell into close contact with whatever is around it (e.g. the substratum or another cell), allowing CglB-mediated substratum-coupling of the Agl–Glt gliding apparatus to then take over and power cell locomotion.
ii. CglD is a cell-surface adhesin that can specifically couple to the Agl–Glt machinery and assist with CglB-mediated substratum-coupling of the complex.

To examine the relationship between CglD and the known components of the gliding apparatus, we first probed the co-occurrence of *cglD* with *cglB* and *gltABCDEFGHIJK* across a range of representative complete bacterial genomes. As with *cglB* and the *glt* genes, *cglD* was highly conserved in members of the order Myxococcales, but was never found to be encoded in clusters for any of the known gliding-apparatus components. Sporadic instances of *cglD* in the absence of most of the gliding-machine genes were also detected, suggesting that *cglD* acquisition may have predated any co-option by the gliding apparatus in the Myxococcales (**fig. S5**).

To further distinguish between the two abovementioned possibilities for the TFM data, we employed bead-force microscopy; therein, using optical tweezers, a large (520 nm-diameter) polystyrene bead was deposited directly on the surface of *M. xanthus* cells, after which CglB-dependent Agl–Glt trafficking events^5,35^ acting on the bead were quantified (**Fig. 6D**). In this manner, a general requirement for CglD to help “recruit” the bead into close contact with the cell surface was negated. Compared to beads placed on WT cells, those deposited on Δ*cglD* cells were trafficked over shorter periods of time and at slower speeds (**Fig. 6E,F**). If CglD did not specifically participate in surface-coupling of the bead to the Glt apparatus, the lengths and speeds of individual bead-run events should have been comparable between WT and Δ*cglD* cells; as this was not the case, the data support a direct involvement of CglD in coupling the internally-trafficked Agl–Glt apparatus to the cell-surface Glt OM platform containing CglB.

### CglD presence is not affected by constituents of the Glt apparatus

As the bead-force microscopy results suggest that CglD functions in concert with the Glt apparatus, and given the previously-demonstrated cellular-retention dependence of CglB on certain constituents of the Glt OM module^5^, we probed for the presence of CglD in mutant strains of all known constituents of the gliding machinery. Unlike CglB^5^, CglD was found to be expressed and retained by the cell independently of any gliding-machinery defect (**Fig. 7A**). Moreover, the folding state of CglD was not affected by the absence of any Glt OM-module protein (**Fig. 7B**), reinforcing its lack of dependence on any Glt components for its retention by cells.

**Fig. 7.**
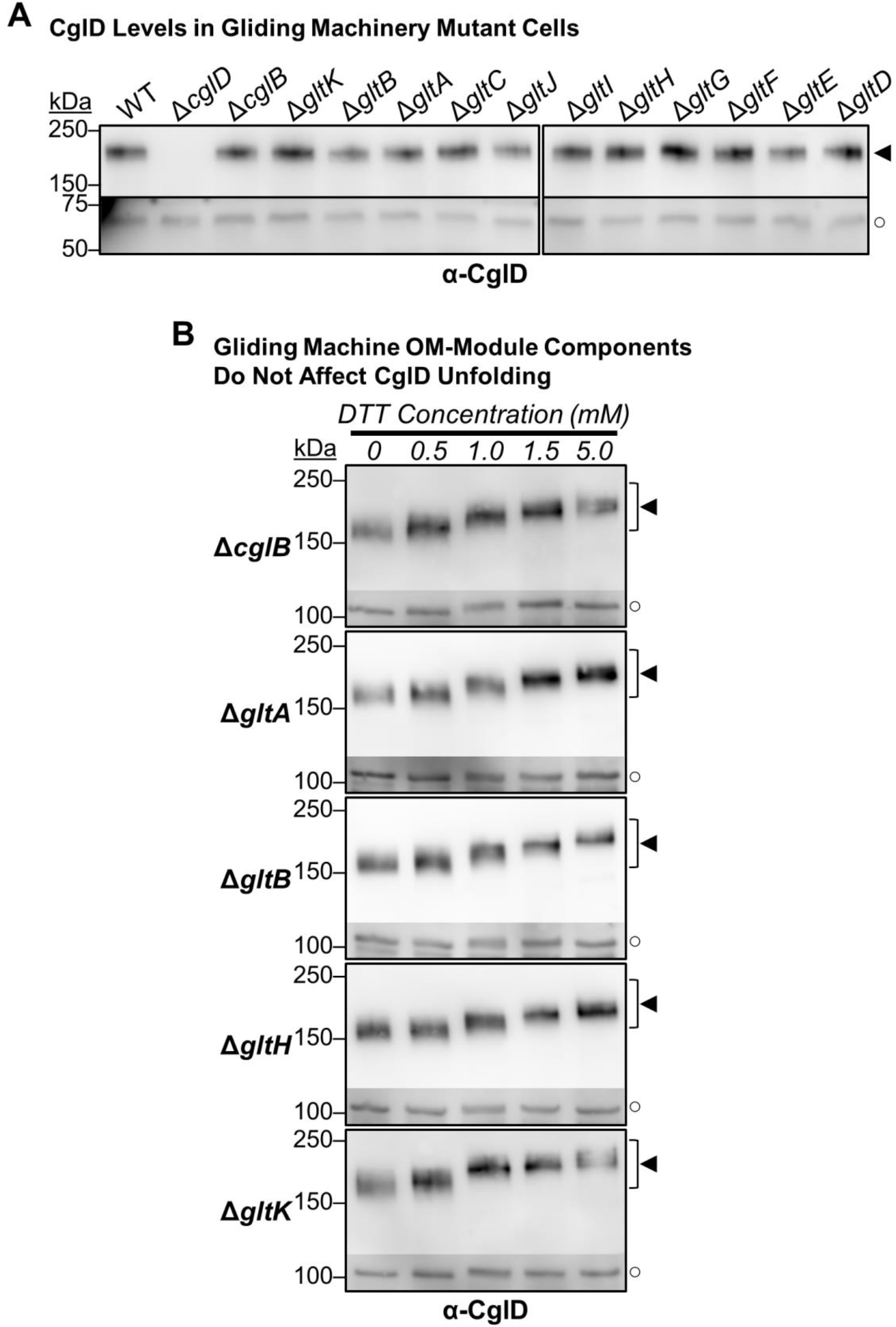
Glt components do not affect the cellular levels and unfolding of CglD. **(A)** α-CglD Western blot of whole-cell extracts from *glt* and *cglB* mutants. The lower, darker zones on the blots corresponds to lower sections of the same blot images for which the contrast has been increased to highlight lower-intensity protein bands. **(B)** α-CglD Western blot of whole-cell extracts from Glt OM-module knockout strains, treated with increasing concentrations of DTT to break disulphide bonds. The lower, darker zones on the blots corresponds to the section of the same blot image for which the contrast has been increased to highlight lower-intensity protein bands. Legend for Panels A and B: ◄, full-length CglD. ○, loading control (non-specific protein band labelled by α-CglD antibody).

### CglD stabilizes bFAs

With the results of bead-force microscopy implicating CglD function as being directly coupled to the gliding machinery, we examined possible structural and functional associations of CglD with the Glt machinery. Using sensitive analyses of raw mass spectra for prey proteins pulled down via purification of glutatathione S transferase (GST)-tagged bait proteins^12^, OM proteins CglD and GltA (as well as periplasmic GltE) were found to co-purify with the GST-tagged C-terminal domain of periplasmic GltD bait (**Table S1**), consistent with a functional linkage for these proteins with the gliding mechanism. Intriguingly, several putative metalloproteases were also pulled down with this bait construct, providing unexpected leads as to potential candidates that may be responsible for cleaving gliding motility adhesins^5^ from the surface of *M. xanthus* cells (**table S1**).

We next probed the effect of CglD absence on the formation of bFAs in gliding cells (**Fig. 8A**). To track the position of the AglRQS-energized Glt trans-envelope apparatus, fluorescently-tagged copies of the AglZ protein are followed in cells^5,6,11,13,35–37^. AglZ is a cytoplasmic filament-forming coiled-coil protein that is required for gliding^36,38^. We thus first compared bFA formation in gliding cells expressing AglZ-YFP with(out) CglD via fluorescence microscopy (**Fig. 8A**). Contrary to the well-defined and compact bFA clusters formed by WT cells, CglD-deficient cells formed larger-yet-less-intense bFA clusters (**Fig. 8B,C**), suggesting that bFAs in the absence of CglD are considerably more diffuse and misshapen. These bFA clusters in Δ*cglD* cells were also more prone to slippage, i.e. slight shifts in anchored position relative to the substratum, than WT cells (**Fig. 8A**), suggesting an inefficient engagement of the bFA with the substratum in the absence of integrin-like CglD.

**Fig. 8.**
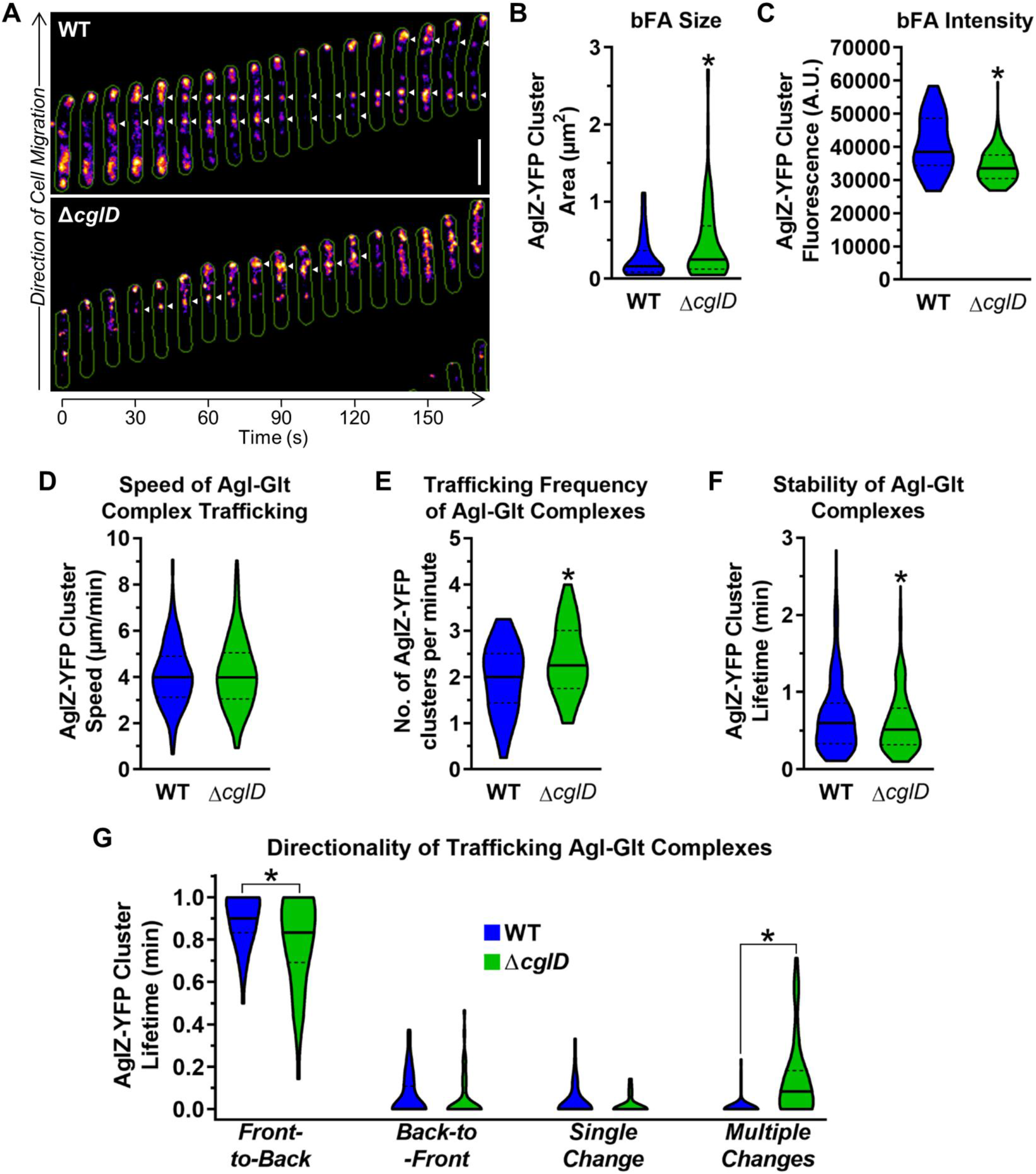
CglD deficiency impacts numerous bFA properties. **(A)** Fluorescence microscopy montage of gliding cells indicating bFA positions (*white arrowheads*) via AglZ-YFP fluorescence. **(B)** Violin plot of bFA cluster size in WT and Δ*cglD* cells (n = 45 and 123, respectively). **(C)** Violin plot of bFA cluster intensity in WT and Δ*cglD* cells (n = 45 and 123, respectively). **(D)** Speed of Agl–Glt complex trafficking via TIRFM (of AglZ-YFP) on chitosan-coated glass surfaces in PDMS microfluidic chambers for WT and Δ*cglD* (n = 259 and 253 clusters, respectively) strains. **(E)** Frequency of trafficking AglZ-YFP complexes via TIRFM (of AglZ-YFP) on chitosan-coated glass surfaces in PDMS microfluidic chambers for WT and Δ*cglD* (n =44 and 43 cells, respectively) strains. **(F)** Stability of trafficking Agl–Glt complexes via TIRFM (of AglZ-YFP) on chitosan-coated glass surfaces in PDMS microfluidic chambers for WT and Δ*cglD* (n = 333 and 346 clusters, respectively) strains. **(G)** Directionality of trafficked Agl–Glt complexes via TIRFM (of AglZ-YFP) on chitosan-coated glass surfaces for WT and Δ*cglD* (n = 44 and 43 cells, respectively) strains. “Front” and “back” are defined as cell poles with high and low AglZ-YFP fluorescence intensity, respectively. For Panels B–G, the lower and upper boundaries of the plots correspond to the minimum and maximum values of the dataset, with the 25th and 75th percentiles displayed (*dashed black lines*). The median (*solid black line*) of each dataset is indicated. Asterisks (*) denote datasets displaying statistically significant differences in distributions (*p* < 0.05) between various strains or conditions, as determined via unpaired two-tailed Mann–Whitney tests.

To achieve high temporal and spatial resolution, we further confined our AglZ-YFP-tracking analysis to clusters along the ventral plane of cells using total internal reflection fluorescence microscopy (TIRFM)^4,5^ (**Fig. S6**). While Glt complexes were found to traffic at equivalent speeds in both WT and Δ*cglD* cells (**Fig. 8D**), complexes in the latter cells were trafficked at a higher frequency (**Fig. 8E**) and were less stable (**Fig. 8F**). Moreover, in the absence of CglD, fewer Glt complexes were found to traffic exclusively from the leading to the lagging cell poles, with more instead demonstrating oscillatory behaviour with multiple changes in trafficking direction (**Fig. 8G**).

Taken together, these data indicate that while CglD is not required for bFA formation, the β-integrin-like protein has a profound impact on the stability of bFA clusters needed for efficient gliding motility in *M. xanthus*.

## Discussion

Based on the data presented in our investigation on *M. xanthus* CglD (from a unicellular bacterium that exhibits true multicellular physiology), this protein possesses the hallmarks of a cell-surface β-integrin-like lipoprotein that is directly involved in gliding motility via bFA stabilization. This finding is supported by multiple lines of evidence:

i. CglD possesses β-integrin-like architecture, including a VWA domain (commonly found in ECM-interacting proteins), and also EGF-like repeats within a Cys-rich stalk
ii. CglD is a surface-localized lipoprotein that co-elutes with members of the gliding motility apparatus
iii. In the absence of CglD, trafficking motility complexes become poorly immobilized, consistent with non-optimal adherence of the complexes to the substratum. Furthermore, without CglD, these trafficking complexes are severely compromised for the transport of surface-associated cargo.
iv. Cells lacking CglD display destabilized bFA clusters that oscillate more frequently, dissociate quicker, and display more diffuse signal patterns.

Below, we discuss potential CglD-modulated gliding complex adhesion as well as future avenues of investigation.

### CglD-like β-integrin proteins

The presence of α/β-integrin-mediated adhesion machinery on the eukaryotic tree of life is an ancient occurrence, pre-dating the appearance of unikonts, i.e. eukaryotic cells with a single flagellum, believed to be the ancestor of all metazoans^39^. The long-standing evolutionary importance of these proteins is consistent with their capacity to bind not only ECM components in metazoans, but also numerous non-ECM ligands^40^.

Integrins were once thought to be exclusive to metazoans, but were later identified in so-called “lower” eukaryotes^39,41^. Exceptionally, the lone detection of a (β) integrin in a prokaryote (cyanobacterium *Trichodesmium erythraeum*) was attributed to a horizontal gene-transfer event^39^. However, from our detection herein of β-integrin-like CglD homologues in a diverse selection of δ-proteobacteria, these data would support an ancestral function/role for β-integrin-like proteins in δ-proteobacterial physiology. It can be envisioned that such proteins could also be involved in adhesion to various substrata and/or other cells, but the link with gliding-motility machinery may only have developed past the divergence of the suborder Cystobacterineae, as the species therein encode CglD in addition to the full complement of known gliding machinery proteins (i.e. GltABCDEFGHIJK+CglB).

### β-integrin-like CglD as a mechanosensor and mechanotransducer for bFA initiation & stabilization

Integrins in eukaryotic cells have long been known to serve as biomechanical sensors of the local environment, being able to distinguish between different substratum rigidities, and in turn transmitting this information via an outside–in mechanism to effect internal cellular changes that regulate adhesion and eukaryotic cell motility^1^. This process is very quick, with integrins being able to detect force and transmit a signal to augment adhesion in under a second^42^. Herein, we have shown that substratum polymer alignment and rigidity is a mitigating factor for CglD-deficient cells, whereas WT cells are more adaptable and highly motile. This would be consistent with β-integrin-like CglD having a role in distinguishing between soft/hard matrices, akin to the scenario with eukaryotic integrins.

As traction forces are applied on nascent eukaryotic integrin adhesions, this leads to integrin clustering and the maturation of eFAs^1^. In the absence of CglD, we observed gliding *M. xanthus* cells to be defective for bFA clustering and stability, with the bFA signal in Δ*cglD* cells being less intense, more diffuse, shorter lived, and highly oscillatory. Thus, in addition to its mechanosensory capacity, our data support a mechanotransductory role for CglD in initiating and maintaining AglRQS directed motorized transport at bFA sites.

### Potential CglD interactions with the gliding apparatus

Mechanosensory and mechanotransductory functions for CglD would heavily implicate interactions of this β-integrin-like protein with the OM module of the trans-envelope gliding apparatus, considering CglD is an OM lipoprotein with no discernible OM-spanning domains. This contention is supported by the analysis of mass spectrometry data from GST-tagged pulldowns in broth-grown cells; through use of a GST-tagged C-terminal domain of periplasmic GltD as bait, periplasmic GltE, but more importantly OM CglD and GltA were reproducibly pulled down. These data could indicate that in a non-surface-engaged state, CglD may already associate with members of the Glt OM module. Functional analogies exist for eukaryotic integrin proteins: when not engaged/activated, integrins are believed to adopt a “resting state” conformation that does not fully associate with their ligands. However, upon binding of the target ligand, integrins undergo a conformational change to an “activated state” in which the VWA domain interacts with the ligand, changing the conformation of the ectodomains into an unbent form. In turn, this transmits a signal across the cytoplasmic membrane to the tail domain of each integrin subunit, leading to activation of multiple cytoplasmic proteins and eFA formation. Interestingly, integrins are receptors involved in cellular adhesion and COMP in humans has been found to affect cellular attachment. Moreover, COMP-based mediation of cell attachment is carried out through interaction with α5β1 integrin in the presence of Ca^2+^ ions^43^. Such COMP–integrin interactions have been proposed to be mediated via COMP binding of the MIDAS motif of β-integrin subunits^44,45^.

As such, the integrated Ca^2+^-binding COMP domain of CglD may promote interaction with an integrin-like protein. As the presence/absence of Ca^2+^ or Glt OM-module proteins did not alter its own conformational stability in broth-grown cells, this may indicate that the CglD COMP domain may be responsible for interactions with the portions of the Glt OM-module, most likely CglB (as the latter is structurally homologous with an integrin αI-domain VWA module)^5^; however, such an interaction would be favoured in a gliding cell in which the substratum has been actively engaged by CglD in concert with CglB-GltABHK. This in turn would result in stabilization of the bFA, allowing the anchored gliding motility complexes accumulated at this site to promote efficient single-cell gliding motility.

### Potential role for the glycocalyx in bFA activity

The glycocalyx of eukaryotic cells has been shown to greatly impact integrin-mediated cell adhesion and force transduction. The mechanical resistance of this (protein-impregnated) cell-surface polysaccharide layer can regulate the clustering of integrins^46,47^. Intriguingly, for almost a century myxobacterial single-cell gliding motility has been associated with a so-called “slime” polysaccharide^48^. Though the phase-bright nature of trails commonly found behind gliding myxobacteria on agar pads was discovered to simply be due to physical depressions left behind in the agar matrix^49^, a polymeric substance left behind gliding cells was still detected on rigid substrata^50^. This deposited polymer is distinct from the known secreted exopolysaccharide (EPS), biosurfactant polysaccharide (BPS), and major spore coat (MASC) polysaccharide, as well as the LPS-capping O-antigen polysaccharide ^8,9,50^ of *M. xanthus*. Moreover, gliding *M. xanthus* cells detected over slime trails were suggested to be more strongly adhered to the substratum^50^. In line with known eukaryotic integrin function and the data presented herein, an important role of slime with respect to *M. xanthus* gliding may thus be to facilitate CglD-mediated bFA clustering to support efficient single-cell gliding motility. A role for the *M. xanthus* glycocalyx in regulating CglD-dependent gliding promotion would also be consistent with our previous data showing that BPS^−^ cells, which have a more compact cell-surface glycocalyx, are hyper-polymertropic^9^; they are exceptionally responsive (in a gliding-dependent manner^5,27^) to mechanical alignment of polymers in compressed substrata, exhibiting preferential swarm expansion along the aligned polymers.

### Conclusion

Ultimately, the findings in this investigation reinforce and highlight the exciting parallels between bFA-mediated prokaryotic gliding and eFA-mediated eukaryotic motility. In turn, this opens possibilities for understanding the evolution of complexity in integrin-mediated cell translocation and its contribution to the development of true multicellular physiology across biological kingdoms.

## Materials and Methods

### Bacterial cell culture

Wild-type^51^ and mutant strains of *M. xanthus* DZ2 (**table S2**) were grown at 32 °C in CYE liquid medium (1% Bacto Casitone Peptone, 0.5% Yeast Extract, 0.1% MgCl_2_, 10mM MOPS [pH 7.4]) with shaking (220 rpm) or on CYE medium solidified with 1.5% (w/v) agar. Cell resuspensions were carried out in TPM buffer (10 mM Tris-HCl [pH 7.6], 8 mM MgSO_4_, 1 mM KH_2_PO_4_).

### Phenotypic Analyses

For all phenotypic analyses, 5 µL of cells resuspended in TPM (at an optical density at 600 nm [OD_600_] of 5.0) were spotted on various substrata. Gliding-flare observations were acquired with an Olympus SZX16 stereoscope equipped with an ILLT base and UC90 4K camera, using CellSens software (Olympus). For gliding-flare analysis, cells in TPM were spotted on CYE 1.5% (w/v) agar, 1.5% (w/v) carrageenan, or 0.6% (w/v) gellan plates and incubated at 32 °C for 7 h. Flares were imaged using the SDF PLAPO 2×PFC objective, with 6.3× zoom, and a fully-open aperture. The illumination wheel was set halfway between the brightfield and empty slots for optimal cross-illumination of the sample. Image acquisition was carried out using linear colour, 50 ms acquisition time, 16.6 dB gain, and a white balance calibrated against an empty zone of the plate.

For T4P-dependent swarm-spreading and fruiting body analysis, images were acquired with an Olympus SZ61 stereoscope with an ILLT base. Swarm spreading was captured at 0.67× zoom, using dark-field illumination, while fruiting bodies were captured at 32× zoom, using full oblique illumination. For swarm-spreading, cells were spotted on CYE 0.5% (w/v) agar plates, while for fruiting body formation, cells were spotted on CF (0.15% casitone [w/v], 10 mM MOPS, 1 mM KH_2_PO_4_, 8 mM MgSO_4_, complemented with 0.02% (NH_4_)_2_SO_4_ and 2% Na_3_C_6_H_5_O_8_), with 1.5% or 0.5% (w/v) agar for phenotype plates. Swarm-spreading and fruiting-body plates were imaged after incubation for 72 h at 32 °C. Swarm diameters were measured using CellSens software.

### Rheometry and cell detachment

A volume (1 mL) of molten CYE 1.5% (w/v) agar was deposited in 35 mm-diameter FluoroDish (World Precision Intruments) as a substratum for *M. xanthus* cells. After solidification of the medium, cells from overnight cultures of WT/Δ*cglD*/Δ*cglB* expressing OMss-mCherry were resuspended in TPM (OD_600_ 5.0), with 5 µL deposited on the agar matrix followed by incubation at 32 °C for 4 h. To facilitate imaging of swarm disintegration under a wide range of dynamic shear forces, we utilized a customized Rheo-Confocal setup where the parallel plate rheometer (Anton Paar MCR302) was assembled on top of a confocal microsystem (Leica SP8)^52^. Glycerol (60%, 1.5 mL) was added to the FluoroDish between the top spinning disc of the rheometer and the swarm. Fluorescence images are captured by the confocal microscope (mCherry detection, laser: 552 nm, 10× magnification, HCX PL APO CS 10×/0.40 dry objective) prior to rotation of the rheometer (i.e. Shear Force: 0), and after each rotation speed increase to follow swarm disintegration. Images at different shear forces were then analysed with Fiji to measure the fluorescent signal, with readings normalized to that detected at “0” shear force for each strain.

### Polymertropism response

Aspect ratio (AR) analyses were performed using previously-described methods^9,27–29^. In brief, *M. xanthus* cells were grown overnight in CYE medium at 28 °C to a density of approximately 5 × 10^8^ cells/mL. Subsequently, the cells were sedimented, (4 000 × *g*, 10 min), resuspended in CYE broth to a density of 5 × 10^9^ cells/mL, and used to inoculate (4 µL) compressed and uncompressed round 85-mm CTTYE (1% casitone [w/v], 2% yeast extract [w/v], 10 mM Tris-HCl [pH 8.0], 1 mM KH_2_PO_4_, 8 mM MgSO_4_) agar plates. To compress the agar, a section of Tygon tubing (outer diameter: 5.56 mm, length: 1 cm) was inserted against the plate wall. The cells on these plates were inoculated at a distance of 43 mm from the inserted tubing. Following incubation at 30 °C for 24, 52, 90, 120, and 144 hours, the perimeters of the colonies were marked at each time interval. The aspect ratio (AR) of each swarm was calculated for each time point by dividing the colony width by the colony height. A round swarm would yield an AR ≈ 1, while a flattened swarm would have an AR > 1. Linear best-fit lines were determined for each replicate dataset, and the AR/time ratio was calculated. Average slope values were calculated for each strain and normalized to the WT strain.

### SDS-PAGE, in-gel fluorescence, and Western immunoblotting

To probe for AglZ-YFP in-gel fluorescence, 10 mL cultures of WT and Δ*cglD* cells expressing AglZ-YFP were grown overnight in CYE broth with shaking (220 rpm) at 32 °C. Cells from these cultures were then sedimented via centrifuge (5000 × *g*, 5 min, room temperature), followed by decanting of the supernatant and resuspension of the cells in 10 mL of TPM via vortex. The OD_600_ of each TPM resuspension was determined, followed by sufficient removal of volume so that resuspension of the removed cells in 500 µL would yield a final OD_600_ of 2.0; these removed cell volumes were thus sedimented as above, resuspended in 500 µL of 1× Laemmli SDS-PAGE sample buffer lacking reducing agent, then heated at 65 °C (30 min). From these samples, 20 µL of equilibrated cell resuspensions (along with 10 µL of pre-stained protein ladder) were loaded on a 10-well 8% polyacrylamide gel, and resolved via SDS-PAGE at 80 V (45 min) then 120 V (75 min). These gels were then scanned with a Typhoon FLA9500 fluorescence scanner (GE Healthcare), using the 473-nm laser to excite AglZ-YFP, and the BPB1 filter (PMT 800) to capture in-gel fluorescence. Bands corresponding to the pre-stained ladder were excited with the 635-nm laser and detected using the LPR filter (PMT 800). Fluorescence intensity of the detected AglZ-YFP bands was obtained using the “plot lanes” function of ImageJ, after which the area under the curve was determined. This signal was normalized to the faster-migrating autofluorescent band in the same lane. These values were subsequently expressed as a percentage of the WT signal for each biological replicate.

To detect CglD from whole-cell lysates, cells were harvested after overnight growth, washed in TPM buffer, and resuspended at an OD_600_) of 1.0 in 1× Laemmli SDS-PAGE sample buffer containing 5% β-mercaptoethanol. Samples were then boiled for 10 min, and 20 µL of each sample were loaded onto 10-well 1-mm 10% acrylamide gels. Sample resolution through gels was conducted in two stages: 45 min at 80 V through the stacking gel, followed by 105 minutes at 120 V through the resolving gel. Resolved samples were subsequently transferred to nitrocellulose membranes via electroblotting at 100 V for 60 min. Membranes were rinsed in Tris-buffered saline (TBS), blocked with TBS containing 5% milk powder (w/v) at 4 °C for 30 min, then incubated overnight with gentle rocking at 4 °C in TBS containing 0.05% Tween 20 (v/v), 5% milk powder (w/v), and a 1:10 000 dilution of the primary pAb α-CglD anti-serum. The next day, blots were washed twice in TBS with 0.05% Tween 20 and then incubated with a goat α-rabbit secondary antibody conjugated to horseradish peroxidase (Biorad) at a 1:5 000 dilution in TBS with 0.05% Tween 20 and 5% milk for 60 min. After two additional washes in TBS with 0.05% Tween 20, the blots were developed using the SuperSignal West Pico chemiluminescence substrate (Thermo) and captured using an Amersham Imager 600 machine.

### Single-cell microscopy analysis

For phase-contrast microscopy on pads, cells were cultured overnight at 32 °C, washed, and resuspended in TPM buffer to an OD_600_ of 2.0. Resuspended cells were then spotted (3 µL) onto a glass coverslip, atop which a pad made of 1.5% agarose (w/v) in TPM buffer was overlaid. Cells were left to associate with the pad for 5 min before imaging at 32 °C using a Zeiss Axio Observer 7 microscope with a 40× objective and an Axiocam 512 camera. For phase-contrast microscopy on chitosan devices, cells were grown under the same conditions but resuspended in TPM buffer containing CaCl_2_ (2 mM). Subsequently, 1 mL of the cell suspension was loaded into the device. After a brief incubation period, the cells were washed with TPM buffer containing CaCl_2_ before imaging. Similar to the previous setup, the cells were allowed to settle for 15 min before imaging at 32 °C as above. Calculation of cell gliding speeds was performed using the MicrobeJ plugin for FIJI^53^, while reversal events (switching of cell gliding direction) were manually counted.

To probe for AglZ-YFP cluster fluorescence on agarose pads, imaging was conducted using an in-house-made aluminum monolithic microscope equipped with a 1.49 NA/100× objective (Nikon). Imaging was performed with an iXon DU 897 electron-multiplying charge-coupled device (EMCCD) camera (Andor Technology), with illumination achieved using a 488-nm diode-pumped solid-state (DPSS) laser (Vortran Stradus). Sample positioning was carried out using a P611 three-axis nanopositioner (Physik Instrument). LabView (National Instruments) was used to program instrument control and integrate control of all components. From these datasets, kymographs were generated using the “Kymograph Builder” function in FIJI. Manual selection of AglZ-YFP clusters was performed, followed by tracking using the MTrackJ FIJI plugin.

To achieve high temporal resolution for real-time AglZ-YFP trafficking, TIRFM was carried out as previously described^4,5^ using chitosan-coated PDMS microfluidic channels. In summary, cells were injected into the chamber and allowed to adhere for 30 min without flow. Any unadhered cells were subsequently removed by manually injecting TPM. TIRFM was performed on the attached cells using an inverted microscope equipped with a 100× oil-immersion Plan-Achromat objective and a closed-loop piezoelectric stage for active autofocus. AglZ-YFP was excited using a 488-nm laser, and the emission was collected by the objective, passed through a dichroic mirror and band-pass filters, and captured by an EMCCD camera. To capture real-time images of the YFP channel, a total of 500 images were taken at a rate of 20 Hz^4,5^.

### Chitosan coating for single-cell analyses

PDMS microfluidic devices were fabricated using a mold. A PDMS mixture was prepared by combining the polymer and crosslinker (in a ratio of 10:1) using the Sylgard 184 Microchem kit. The mixture was thoroughly mixed and then centrifuged for 5 min (500 × *g*) to remove any trapped air bubbles. The prepared PDMS mixture was then poured onto the mold. The mold was placed under vacuum for 20 min to remove any remaining micro air bubbles from the mixture. Afterward, the mold with the PDMS was incubated in an oven at 65 °C for 2 h. Once the PDMS device was cured, it was carefully separated from the mold, and small holes with a diameter of 1.2 mm were created as entry and exit points. The PDMS devices were then cleaned using ethanol and water. Glass coverslips (for mounting the PDMS devices) were cleaned using the same method and were plasma-activated for 30 min using the Basic Plasma Cleaner (Harrick Plasma) on the “HI” setting. The PDMS devices were also plasma-activated for 2 min on the “LOW” setting. Following activation, the coverslips and PDMS devices were carefully aligned and pressed together. The assembled device was then incubated overnight at 65 °C, then stored at room temperature until needed. Prior to seeding the device with cells, chitosan solution (100 mg chitosan powder [shrimp, <75% deacetylated, Sigma], dissolved in 10 mL dilute acetic acid, pH 4.0) containing increasing concentrations of CaCl_2_ was injected into the channel(s) to be used, allowed to adsorb for 30 min, then washed with TPM (1 mL).

To compare the effect of Ca^2+^ on gliding in the presence/absence of CglD, borosilicate glass microscope cover slips (75 × 25 × 0.17 mm) were first rinsed with 95% EtOH and ddH_2_O, dried under a stream of N_2_ gas, then treated in a plasma cleaner for 10 min to generate silanol groups on the glass surface to improve chitosan adsorption. Each coverslip was then fixed atop a spin-coating pedestal with double-sided tape, after which chitosan solutions (100 mg chitosan dissolved in 10 mL dilute acetic acid, pH 4.0) supplemented with 0, 0.5, or 2.0 mM CaCl_2_ were spotted atop the pedestal-mounted coverslip, followed by spinning at 2000 rpm for 5 min. Coverslips were then carefully removed from the pedestal and dipped into dilute acetic acid solution (pH 4.0) using tweezers to remove all excess deposited chitosan, leaving behind only chitosan chains directly adsorbed to the glass surface, then stored at room temperature. Prior to inoculation with cells, 1 mL of ddH_2_O was added to each coverslip to rehydrate the chitosan. After 30 min, excess water was removed via decanting, then 5 µL of cells resuspended in TPM (OD_600_ 0.5) were added to the centre of each rehydrated coverslip, then covered with a square coverslip and left to adhere for a minimum of 2 h at room temperature. Cells were then imaged on a Nikon Eclipse TE2000 microscope (40× magnification, DIA illumination) at 32 °C for 5 min, with images acquired at 30-s intervals.

### Testing of CglD susceptibility to Proteinase K and DTT

Susceptibility to Proteinase K was assayed as previously described^5^. In brief, cells from overnight cultures were resuspended to an OD_600_ of 2.0 in 600 µL of TPM buffer. Afterwards, 6 µL of Proteinase K (New England Biolabs) was added to the cell resuspension. At each designated time point, 100 µL of digestion reaction volume were removed, mixed with trichloroacetic acid (10% final concentration) to halt digestion, and kept on ice. Digestion aliquots were then twice resuspended in 1 mL acetone and sedimented at 16 000 × *g* (5 min), then left uncapped overnight in a fumehood to allow for residuel acetone to evaporate. Precipitated protein pellets were then resuspended in 100 µL of 1× Laemmli buffer with β-mercaptoethanol. Samples were subsequently boiled and processed for Western blot analysis as outlined above.

To probe for disulfide-based denaturation differences dependent on Glt OM-complex mutant background, Samples were prepared by growing overnight cultures and resuspending cells in TPM buffer at an OD_600_ of 4.0 with various concentrations of DTT, ranging from 0 to 5 mM. The samples were then mixed with 2× Laemmli buffer lacking reducing agent to reach a final OD_600_ of 2.0, followed by boiling and processing for Western blot analysis as outlined above.

### Phylogeny and gene co-occurrence

Sixty-one-order Myxococcales genomes, belonging to three suborders and nine families^54–67^, in addition to 59 outgroup genomes (members from 32 non-Myxococcales Deltaproteobacteria, 4 α-, 6 β-, 9 γ-, 4 ε-proteobacteria, 2 Firmicutes, 1 Actinobacteria, and 1 FCB group organism) were selected for this study. To build a maximum-likelihood phylogenetic tree, gapless concatenated alignment of 26 housekeeping protein sequences^64,68^ was subjected to RAxML with JTT Substitution Model and 100 bootstrap values^69^. Sequential distribution of gliding motility genes, i.e. *agl*, *glt* (M1, G1 and G2 clusters)^11^ along with *cglB*^15,70,71^ and *cglD* was identified within all 120 genomes under study using homology searching via tBLASTn and JackHMMER (HMMER 3.3.2 suite released in Nov. 2020)^72^ with two iterative search rounds and an E-value cut-off of 1e^-5^. Visualization of the relative distribution of gliding motility proteins in the multi-protein phylogeny was done using iTol v6.5.3^73^. The strip to the right of the phylogeny depicts the taxonomic classes (from top to bottom: Myxococcales, non-Myxococcales δ-proteobacteria, α-, β-, γ-, ε-proteobacteria, Actinobacteria, Firmicutes, and Fibrobacteres, respectively).

### Tertiary structure homology detection and protein modeling

Structural homologues to CglD (MXAN_0962) were identified via fold-recognition analysis by HHpred^74^ against protein structures in the PDB_mmCIF70 and PDB_mmCIF30 databases of structures deposited in the Protein Data Bank. Top hits were based on the highest probability scores. Tertiary structure modelling of CglD was carried out using the ColabFold pipeline to run AlphaFold with default settings^75,76^. The highest-confidence CglD model structure was used to generate structural alignments with known proteins using TM-align^77^. All structure figures were created with PyMol.

### Traction force microscopy

To grow the various strains tested, cells were first recovered from a frozen stock via streaking on a CTTYE (1% peptone, 0.2% yeast extract, 10 mM Tris, 1 mM KH_2_PO_4_, and 8 mM MgSO_4_, pH 7.6) 1.5% agar plate, incubated at 32 °C. Vegetative cells were then used to inoculate a 10 mL CTTYE liquid culture in a flask, with shaking incubation overnight at 32 °C.

For TFM imaging, the samples were prepared using 35 mm diameter Petri dishes with a glass bottom (Thermo Fisher, prod. #150682). The inner surface of the glass was plasma cleaned, treated with 2 vol.% 3-(Trimethoxysilyl)propyl methacrylate (TMSPMA) for 2 min, washed three times with pure ethanol, and dried^78^. To generate the polyacrylamide (PAA) hydrogel substrate, a 0.25 mL PAA solution (18.8 μL 40% acrylamide, 7.5 μL 2% Bis, 0.222 mL DI water, 1.25 μL 10% ammonium persulfate solution, and 0.375 μL TEMED) was prepared. Suspended fluorescent particles (5 μL, FluoSpheres Carboxylate-Modified Microspheres, 0.04 µm, red-orange fluorescent (565/580), 5% solids) were also added to the mixture. For each substratum, 15 μL of the PAA solution was dispensed on the glass bottom of the petri dish, overlaid with a 12 mm diameter glass coverslip (Thermo Fisher 12CIR-1), and allowed to gelate for 30 min. Following gelation, the coverslip was gently removed and the substratum was soaked in chitosan solution (10 mg chitosan, dissolved in 3 mL 0.2 M acetic acid, and then diluted 1:50) for at least 45 min. The substratum was then washed three times by adding CTTYE and soaking it for ∼10 min. Finally, all liquid was aspirated from the Petri dish, with excess residue carefully wicked away with a kimwipe.

Upon preparation of the substratum, 2 μL of *M. xanthus* cell suspension (OD_550_ 0.7) was added to the top of the gel matrix, with cells left to adhere for 10 min, followed by removal of excess liquid on top of the gel via wicking with a Kimwipe. Petri dishes were then covered again and incubated at 32 °C for 1 h. Following incubation, a 12 mm diameter coverslip was added on top of the gel and gently compressed so that it was uniformly attached to the gel surface. A chamber was then created around the PAA gel in each Petri dish using a 2 mm-thick laser-cut acrylic spacer and a 22 mm × 22 mm glass coverslip. Lastly, the edges of this chamber were sealed with Valap (1/3 vaseline, 1/3 lanolin, and 1/3 paraffin by weight).

Images for TFM were captured with a commercial Nikon Ti-E microscope with Perfect Focus System (PFS) and Yokogawa CSU-21 spinning disk confocal. We used a Nikon 60× Apo Water Immersion objective with long working distance and an Andor Zyla 4.2 sCMOS camera. There was an extra 1.5× magnification through the base of the microscope, so the total magnification of our images was 90×. In each acquisition, one brightfield image was recorded to observe the cells, followed immediately by a fluorescence acquisition using one laser (561 nm light) for the fluorescent particles.

To prevent damage to the bacterial cells from the laser, laser power was kept low (10%). The time between sequential acquisitions was 15 s. Throughout the imaging process, the samples were kept at 25 °C through use of a temperature-controlled cabinet.

Brightfield and laser images were analyzed separately. For the laser images, the slow drift was first removed via tracking the motion of the fluorescent particles in the substratum and measuring the mean velocity of the substratum as an intact solid body. This was followed by use of a band-pass filter to highlight the fluorescent particles. A custom PIV algorithm was then used to measure the displacement of the particles. Lastly, following established methods^79,80^, traction was reconstructed from the displacement field of the gel. The same analysis method was used for analyzing both the dilute-cell and cell-layer samples.

The brightfield images were used to track the motion of individual cells in the dilute regime. The original images were processed, binarized, and segregated, so that the center of mass for each cell could be located. Cell motion was then tracked using the Matlab version of the Particle Tracking Velocimetry (PTV) code developed by Blair and Dufresne^81^. Using the positions of the cells in every frame, we calculated their speeds. Through analysis of the defined rectangular area (14.7 μm wide and 22 μm long, with the long axis parallel to the cell body) centered around individual cells, the distribution of traction generated by single cells was measured. For cell layers, the motion of individual cells was no longer tracked. Regions for analyses were selected in which the cells were concentrated and formed a monolayer, with traction measurements recorded across the entire field of view.

### Flow chamber construction and bead-force microscopy

For bead-force microscopy assays, AglZ-YFP-expressing WT and Δ*cglB* strains of *M. xanthus* were grown shaking in CYE broth overnight at 32 °C to OD_600_ ∼0.6., after which 1 mL of culture was sedimented (8000 rpm, 5 min). The pellet was resuspended in 400 µL TPM buffer. Flow chambers were constructed using two layers of double-sided tape, a 1 mm-thick microscope slide, and a 100 μm-thick glass cover slip (#1.5) to allow a final volume of approximately 60 μL as previously described^5,35^. To facilitate cell attachment, agarose (40 μL at 0.7% w/v) dissolved in 6 M DMSO was injected into the chamber and allowed to sit at room temperature for 15 min. The chamber was then washed with 400 μL TPM, followed by injection of *M. xanthus* cells (60 μL) into the chamber and left to sit at room temperature onto the agarose-coated surface for 30 min. Unattached cells were then thoroughly washed away with a total of 2 mL TPM media containing 10 mM glucose. The flow chamber was then mounted onto the microscope for imaging.

Uncoated polystyrene beads (diameter 520 nm; Bangs Laboratories) were washed and diluted in 1 mL TPM containing 10 mM glucose and injected into the chamber (1 μL). Single beads were optically trapped and placed near the midpoint of the cell length for each immobilized cell of interest.

### Bead tracking and video analysis

Cells of *M. xanthus* with surface-deposited beads were imaged for 3 min, with images captured every 10 s. Movies were analyzed using a custom MATLAB tracking code. Prior to tracking beads, the code filtered and subtracted the background of the image from the cell-attached bead. An internal MATLAB centroid function was then used to identify the center of the bead and converted the *x*,*y* pixel values of the center of the bead in each frame to microns. The *x*,*y* position values of the bead center were then used to compute bead displacements to identify and extract individual motor-driven bead runs. Injection of nigericin (20 µM), a pH gradient/proton-motive force inhibitory drug, into the flow chamber with WT cells was also carried out; this drug was used to disable the molecular motors and reduce bead motion without impacting motor force production allowing us to determine a threshold for a bead run. Similar previous experiments led to negligible bead motion^35^. Bead runs were characterized as how far the bead displaced above the determined threshold in a single given direction without halting. The displacement for each individual motor-driven bead run was used to compute the average bead speed (µm/min) per run for WT and Δ*cglD* cells.

### Analysis of GST affinity chromatography data via mass spectrometry

In duplicate, C-terminal amino acids 800–1218 of *M. xanthus* GltD (formerly AgmU), fused to glutathione-S-transferase (GST), were previously heterologously expressed in *E. coli*, purified, and used as bait to pull down potential interactors from *M. xanthus* whole-cell lysate (with GST by itself used as a control to identify non-specific binders). Pulldown samples were then digested with trypsin, with raw tandem mass spectra acquired at the UC Berkeley Proteomics/Mass Spectrometry Laboratory using a Thermo LTQ XL mass spectrometer^12^. Herein, the raw MudPIT mass spectra from these GST-affinity pulldowns were processed using Thermo Proteome Discoverer software (v2.4.1.15) with the SEQUEST search engine at the Concordia University Centre for Biological Applications of Mass Spectrometry. Database searches were carried out against the UniProt *Myxococcus xanthus* DK 1622 proteome database (UP000002402, v2017-10-25). The enzyme for the database search was chosen as trypsin (full), with maximum missed cleavage sites set to 3. Mass tolerances of the precursor and fragment ions were set at 1.0 Da. Dynamic modifications on Methionine (oxidation, +15.995 Da), protein N-terminus (Acetyl, +42.011; Met-loss, −131.040; Met-loss-Acetyl, −89.030) and static modifications on Cysteine (carbamidomethyl, +57.021 Da) were allowed. Only peptides with high confidence were reported. The XCorr confidence thresholds were applied with the factory default values, as 1.5 for z = 1, 2.0 for z = 2, 2.5 for z = 3 and 3.0 for z >=4 ions. To stringently identify “MXAN_” proteins pulled down via the GltD bait, non-specific hits pulled down with the GST-alone controls were first subtracted from GltD-GST hit lists, followed by removal of hits not meeting the quality threshold (2.5 minimum peptides, 26 average score, 16% average coverage) and not detected across both replicates.

### Statistical analysis

For all comparisons of dataset distributions (**Figs. 3D, 4B,C, 5A,B, 6A,C,E,F, 8B-G**) analyses of statistical significance were carried out via unpaired two-tailed Mann-Whitney test. Differences in mean values for swarm diameter were evaluated for statistical significance using a Wilcoxon signed-rank test performed relative to the reference value of “100” for WT samples (**Fig. 3B**). All statistical analyses were carried out in GraphPad Prism (version 8) at a confidence interval of 95% (*P < 0.05*).

## Acknowledgments

The authors would like to thank A. Roussel and R. Vincentelli (CNRS–Aix-Marseille University, Architecture et fonction des macromolécules biologiques) for CglD protein used to generate pAb; the Confocal Imaging Facility, a Nikon Center of Excellence, in the Department of Molecular Biology at Princeton University for instrument use and technical advice; G. Laevsky in particular for his support and suggestions.

## Funding

Natural Sciences and Engineering Research Council of Canada, Discovery grant RGPIN-2016-06637 (STI)

Natural Sciences and Engineering Research Council of Canada, Discovery grant RGPIN-2023-05576 (STI)

Natural Sciences and Engineering Research Council of Canada, Discovery grant RGPIN-2020-07169 (AJE)

Natural Sciences and Engineering Research Council of Canada, grant EQPEQ/472339-2015 (AJE)

Banting Research Foundation, Discovery Award 2018-1400 (STI)

Canadian Institutes of Health Research postdoctoral fellowship (STI)

Aix-Marseille University AMIDEX Excellence Program (STI)

PROTEO, The Quebec Network for Research on Protein Function, Engineering, and Applications studentship (NYJ, FS)

European Research Council, Advanced Grant JAWS (TM)

Bettencourt-Schueller Foundation, Coup d’élan pour la recherche française 2011 (TM)

National Science Foundation grant CAREER PHY-0844466 (JWS)

National Science Foundation grant PHY-1734030 (EH) Fondation ARC studentship (LMF)

Glenn Centers for Aging Research award (BPB)

National Institutes of Health award P50 GM071508 (BPS)

National Institutes of Health grant GM20509 (DRZ)

National Institutes of Health grant GM129000 (BN)

Indian Institute of Technology Hyderabad seed grant (GS)

Department of Science and Technology, India INSPIRE grant (GS)

## Author contributions

Conceptualization: NYJ, STI

Methodology: NYJ, EH, AB, NK, DJL, HJ, GS, JBF, MD, STI

Software: EH, AMB

Investigation: NYJ, EH, AB, FS, NK, DJL, LMF, BF, UM, JBF, BPB, MD, BN, STI

Visualization: NYJ, EH, STI

Supervision: GS, DRZ, GS, AG, MN, AJE, OT, JWS, TM, STI

Writing—original draft: NYJ, STI

Writing—review & editing: NYJ, EH, AB, GS, TM, STI

## Data and materials availability

All data are available in the main text or the supplementary materials.

