## Supplementary material for "Integrin-like adhesin CglD confers traction and stabilizes bacterial focal adhesions involved in myxobacterial gliding motility": Figs. S1 to S6, Table S2

**This PDF file includes:**

Figs. S1 to S6
Table S2

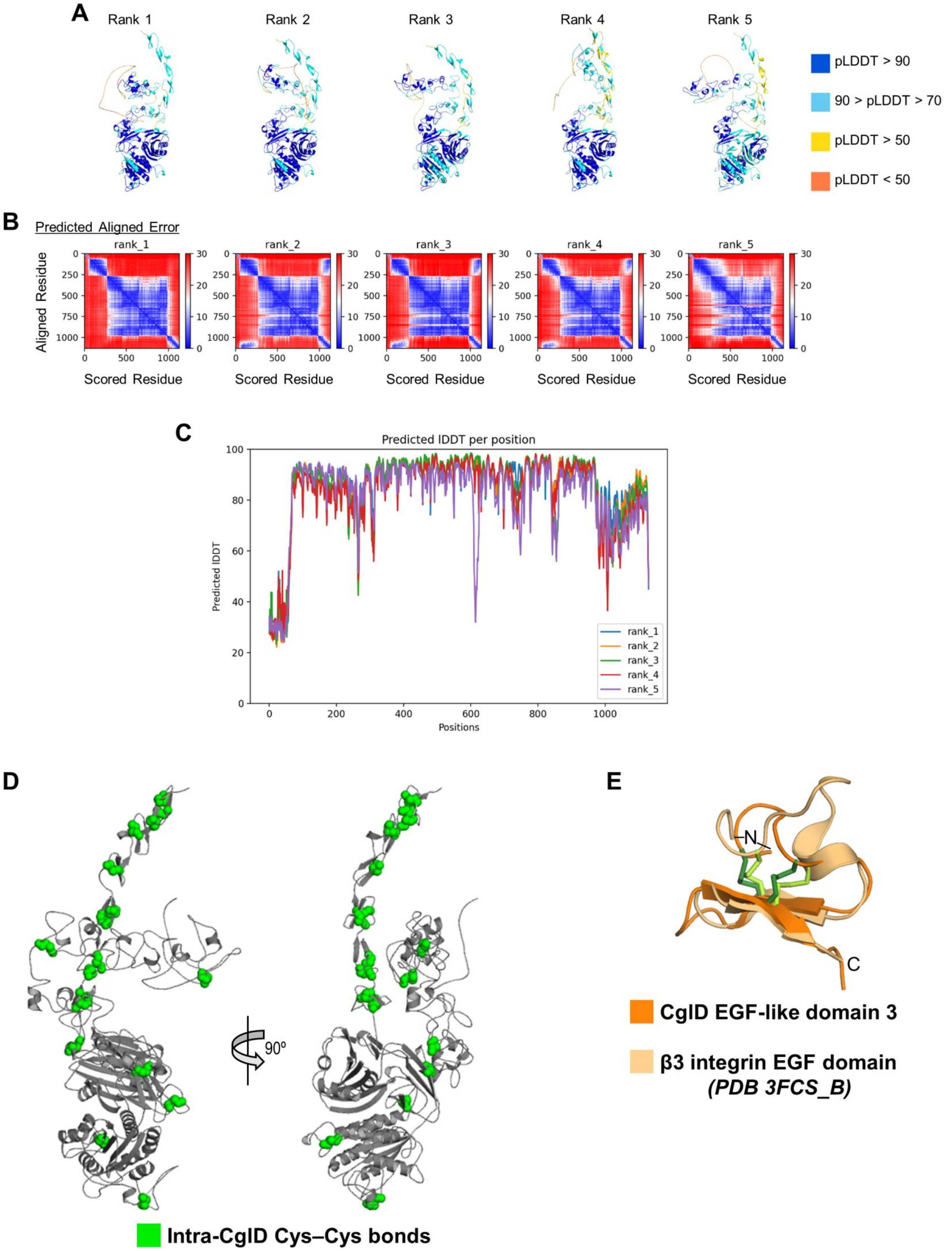
A
Rank 1
Rank 2
Rank 3
Rank 4
Rank 5
pLDDT > 90
90 > pLDDT > 70
pLDDT > 50
pLDDT < 50
B
Predicted Aligned Error
rank_1
rank_2
rank_3
rank_4
rank_5
Aligned Residue
Scored Residue
C
Predicted lDDT per position
Predicted lDDT
Positions
rank_1
rank_2
rank_3
rank_4
rank_5
D
90º
Intra-CglD Cys–Cys bonds
E
N
C
CglD EGF-like domain 3
β3 integrin EGF domain
(PDB 3FCS_B)

**Fig. S1. CglD AlphaFold prediction quality and structural homology.**

**(A)** AlphaFold CglD models ranked from 1 to 5. Per-residue confidence scores (pLDDT: predicted local-distance difference test) are directly reported on the 3D structure of CglD (with the range of possible values from <50 [*orange*] to >90 [*blue*]).
**(B)** Corresponding predicted aligned error for each ranked model.
**(C)** Plot showing the corresponding pLDDT scores per residue for each ranked model.
**(D)** CglD model structure with highlighted (*green balls*) disulfide (Cys–Cys) bonds.
**(E)** Structural alignment of EGF-like Domain 3 of the CglD model structure (*dark orange*) with the EGF domain from β3-integrin (PDB: 3FCS_B) (*pale orange*). Disulfide bonds within each structure are indicated in *dark green* and *pale green*, respectively. Structures were aligned using TMalign.

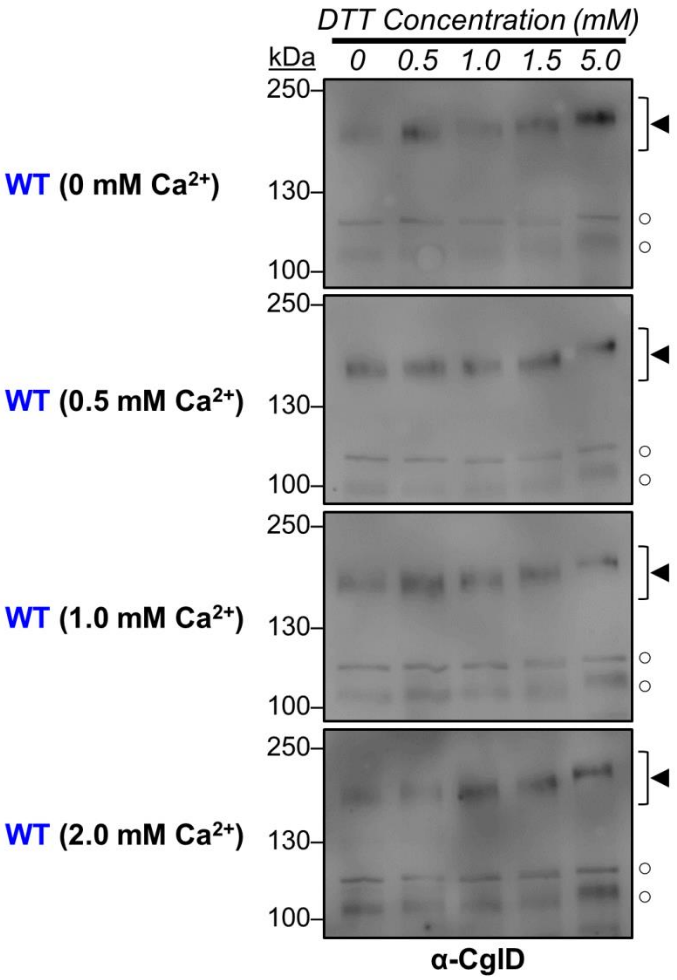


**Fig. S2. $Ca^{2+}$ effect on CglD unfolding.**

α-CglD Western blots of WT whole-cell extracts treated with increasing concentrations of DTT to break disulphide bonds. The four blots correspond to WT cultures grown in various concentrations of $CaCl_2$ (0 to 2 mM). Legend: ◄, full-length CglD; ○, loading controls (labelled non-specifically by α-CglD pAb).

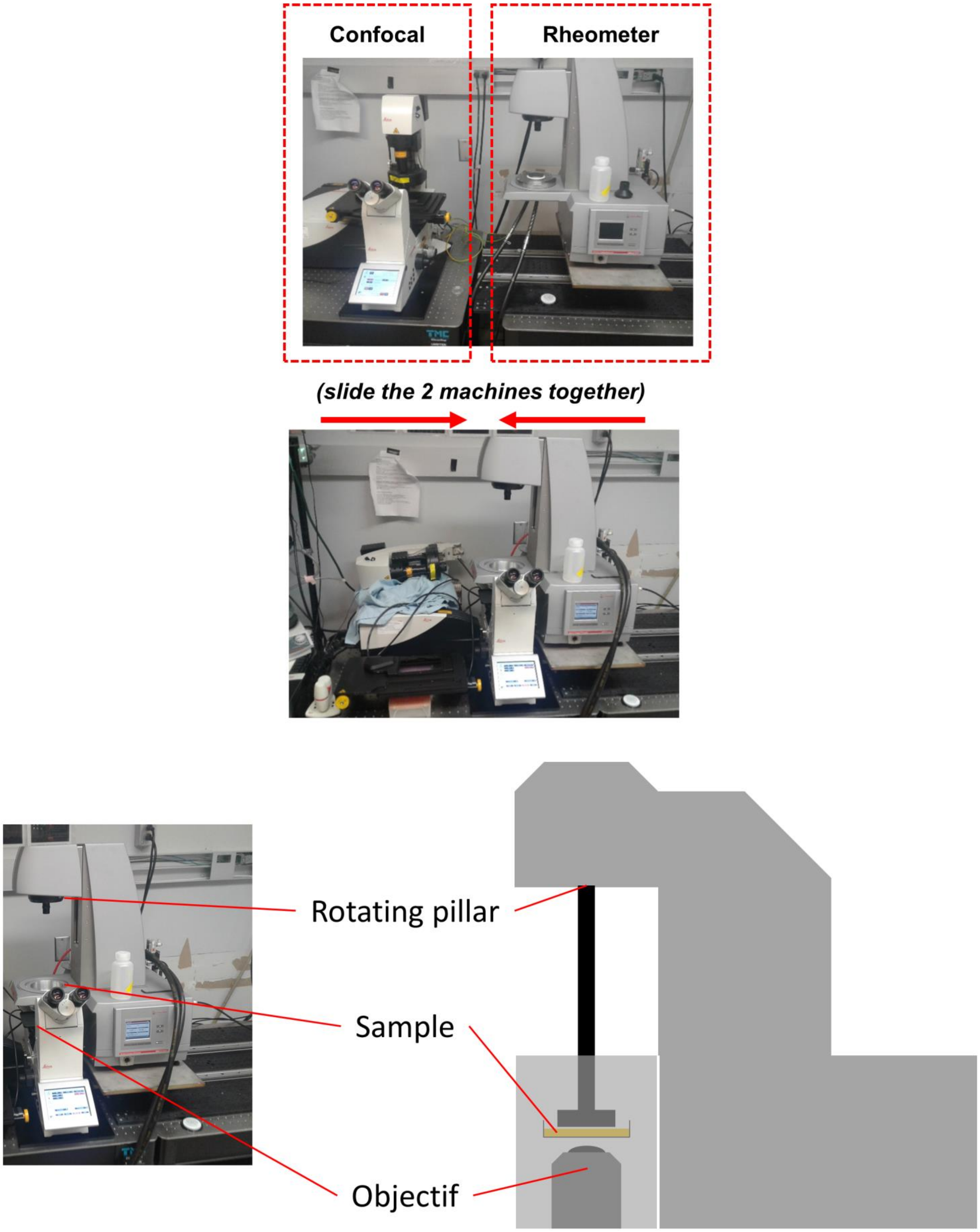


**Fig. S3. Depiction of the combined microscope and rheometer setup.**

Top: Photographs of the separated microscope (left) and rheometer (right) (top pictures). Bottom: Photograph of the combined setup used for the rheometry imaging experiments. The diagram represents the combined device depicted with a sample on the glass-bottomed agar-coated fluorodishes used for imaging.

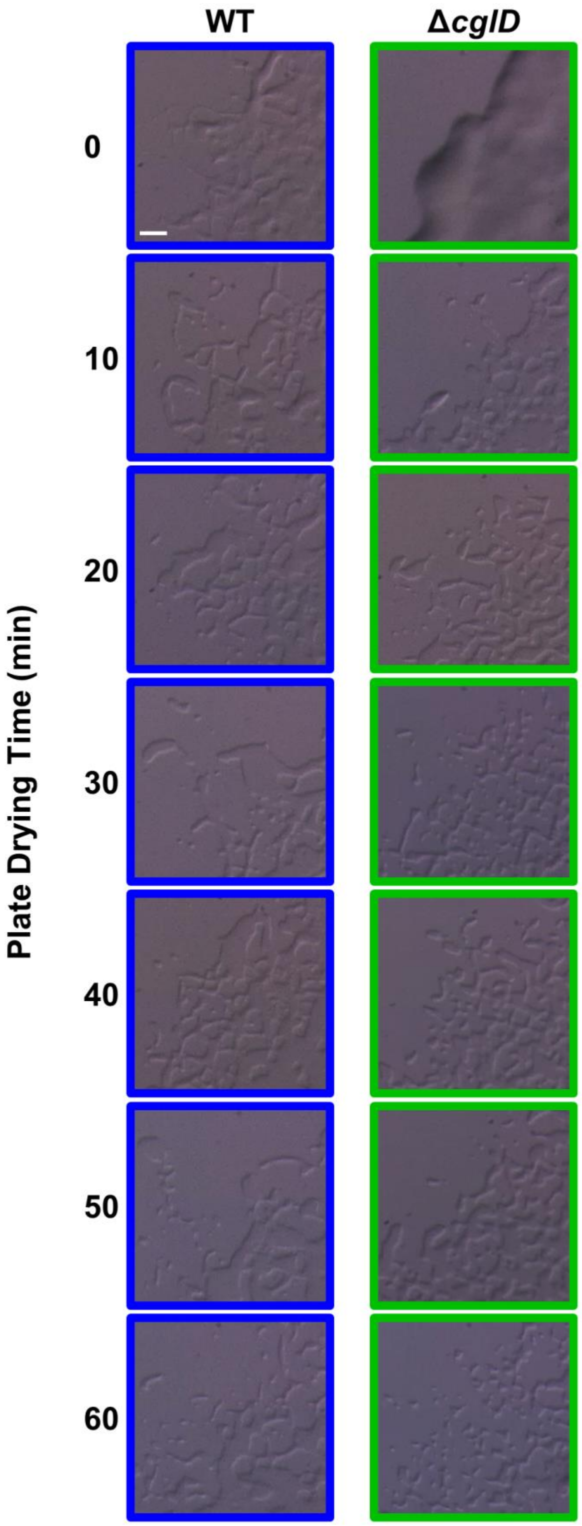


**Fig. S4. CglD sensitivity to the dryness of the substrata.**

Gliding motility flares on CYE hard substrata (scale bar: 50 µm) for WT (*blue*) and Δ*cglD* (*green*) cells. Each time point corresponds to the time that the CYE 0.5% agar plate was left to dry uncovered under the biohood prior to 2 h of (covered) drying on the benchtop at room temperature, after which the cell suspensions were spotted. Each image was captured after 24 h at 32 °C.

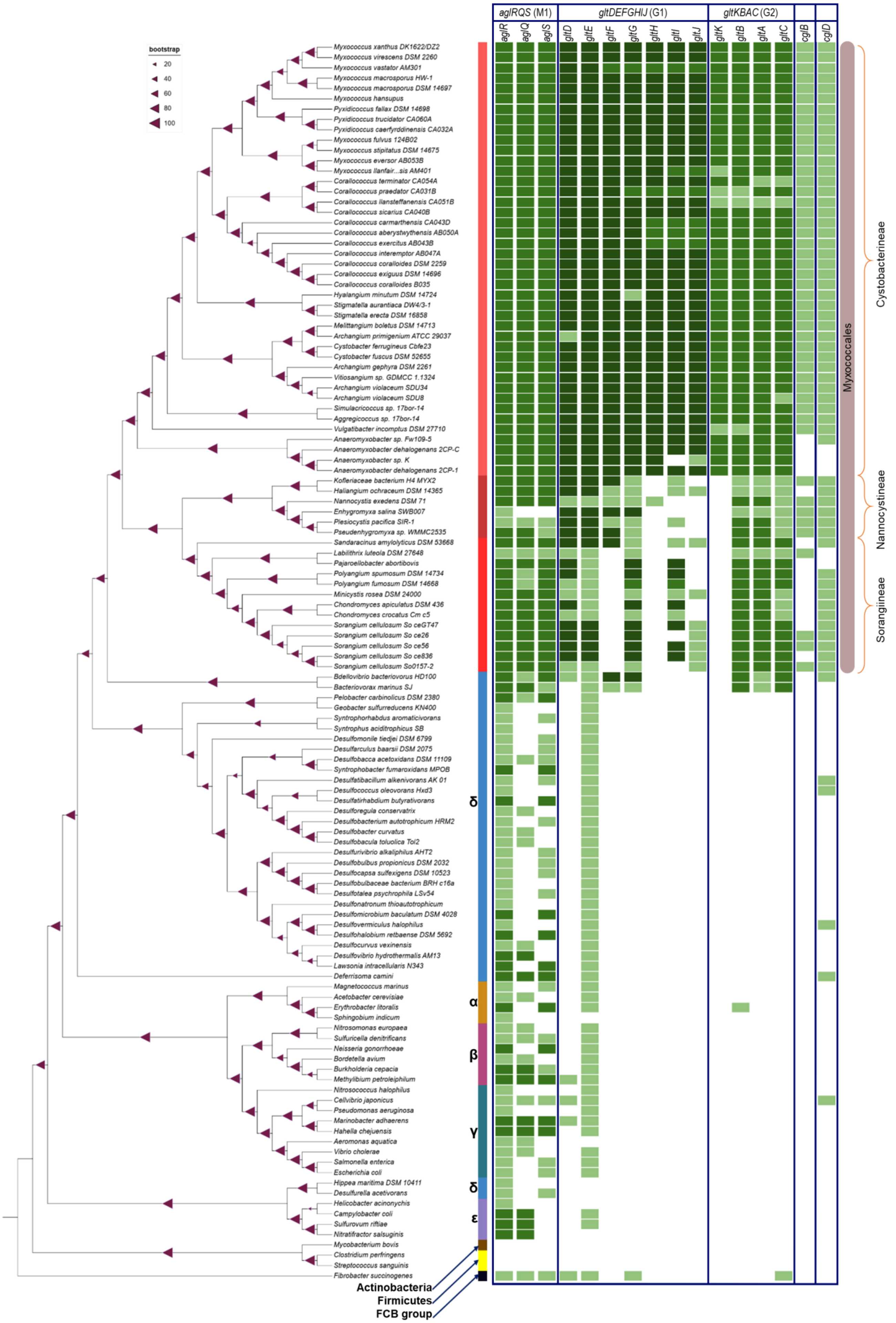

aglRQS (M1)
gltDEFGHIJ (G1)
gltKBAC (G2)
aglR
aglQ
aglS
gltD
gltE
gltF
gltG
gltH
gltI
gltJ
gltK
gltB
gltA
gltC
cglB
cglD
bootstrap
20
40
60
80
100
Myxococcus xanthus DK1622/DZ2
Myxococcus virescens DSM 2260
Myxococcus vastator AM301
Myxococcus macrosporus HW-1
Myxococcus macrosporus DSM 14697
Myxococcus hansupus
Pyxidicoccus fallax DSM 14698
Pyxidicoccus trucidator CA060A
Pyxidicoccus caerfyrddinensis CA032A
Myxococcus fulvus 124B02
Myxococcus stipitatus DSM 14675
Myxococcus eversor AB053B
Myxococcus llanfair...sis AM401
Corallococcus terminator CA054A
Corallococcus praedator CA031B
Corallococcus llansteffanensis CA051B
Corallococcus sicarius CA040B
Corallococcus carmarthensis CA043D
Corallococcus aberystwythensis AB050A
Corallococcus exercitus AB043B
Corallococcus interemptor AB047A
Corallococcus coralloides DSM 2259
Corallococcus exiguus DSM 14696
Corallococcus coralloides B035
Hyalangium minutum DSM 14724
Stigmatella aurantiaca DW4/3-1
Stigmatella erecta DSM 16858
Melittangium boletus DSM 14713
Archangium primigenium ATCC 29037
Cystobacter ferrugineus Cbfe23
Cystobacter fuscus DSM 52655
Archangium gephyra DSM 2261
Vitiosangium sp. GDMCC 1.1324
Archangium violaceum SDU34
Archangium violaceum SDU8
Simulacricoccus sp. 17bor-14
Aggregicoccus sp. 17bor-14
Vulgatibacter incomptus DSM 27710
Anaeromyxobacter sp. Fw109-5
Anaeromyxobacter dehalogenans 2CP-C
Anaeromyxobacter sp. K
Anaeromyxobacter dehalogenans 2CP-1
Kofleriaceae bacterium H4 MYX2
Haliangium ochraceum DSM 14365
Nannocystis exedens DSM 71
Enhygromyxa salina SWB007
Plesiocystis pacifica SIR-1
Pseudenhygromyxa sp. WMMC2535
Sandaracinus amylolyticus DSM 53668
Labilithrix luteola DSM 27648
Pajaroellobacter abortibovis
Polyangium spumosum DSM 14734
Polyangium fumosum DSM 14668
Minicystis rosea DSM 24000
Chondromyces apiculatus DSM 436
Chondromyces crocatus Cm c5
Sorangium cellulosum So ceGT47
Sorangium cellulosum So ce26
Sorangium cellulosum So ce56
Sorangium cellulosum So ce836
Sorangium cellulosum So0157-2
Bdellovibrio bacteriovorus HD100
Bacteriovorax marinus SJ
Pelobacter carbinolicus DSM 2380
Geobacter sulfurreducens KN400
Syntrophorhabdus aromaticivorans
Syntrophus aciditrophicus SB
Desulfomonile tiedjei DSM 6799
Desulfarculus baarsii DSM 2075
Desulfobacca acetoxidans DSM 11109
Syntrophobacter fumaroxidans MPOB
Desulfatibacillum alkenivorans AK 01
Desulfococcus oleovorans Hxd3
Desulfatirhabdium butyrativorans
Desulforegula conservatrix
Desulfobacterium autotrophicum HRM2
Desulfobacter curvatus
Desulfobacula toluolica Tol2
Desulfurivibrio alkaliphilus AHT2
Desulfobulbus propionicus DSM 2032
Desulfocapsa sulfexigens DSM 10523
Desulfobulbaceae bacterium BRH c16a
Desulfotalea psychrophila LSv54
Desulfonatronum thioautotrophicum
Desulfomicrobium baculatum DSM 4028
Desulfovermiculus halophilus
Desulfohalobium retbaense DSM 5692
Desulfocurvus vexinensis
Desulfovibrio hydrothermalis AM13
Lawsonia intracellularis N343
Deferrisoma camini
Magnetococcus marinus
Acetobacter cerevisiae
Erythrobacter litoralis
Sphingobium indicum
Nitrosomonas europaea
Sulfuricella denitrificans
Neisseria gonorrhoeae
Bordetella avium
Burkholderia cepacia
Methylibium petroleiphilum
Nitrosococcus halophilus
Cellvibrio japonicus
Pseudomonas aeruginosa
Marinobacter adhaerens
Hahella chejuensis
Aeromonas aquatica
Vibrio cholerae
Salmonella enterica
Escherichia coli
Hippea maritima DSM 10411
Desulfurella acetivorans
Helicobacter acinonychis
Campylobacter coli
Sulfurovum riftiae
Nitratifractor salsuginis
Mycobacterium bovis
Clostridium perfringens
Streptococcus sanguinis
Fibrobacter succinogenes
δ
α
β
γ
δ
ε
Actinobacteria
Firmicutes
FCB group
Myxococcales
Cystobacterineae
Nannocystineae
Sorangiineae

**Fig. S5. Co-occurrence and gene synteny studies of *cglD* in bacteria with a focus on Proteobacteria.**

Taxonomic distribution and co-occurrence of *agl*, *glt*, *cglB*, and *cglD* genes in Proteobacteria. Bootstrap values at each node are indicated as shown in the left-side legend. Colour of gene hit indicates synteny with the G1 *gltDEFGHIJ* (*dark green*) or G2 *gltKBAC* (*green*) gene clusters or lack thereof (*light green*), respectively. Herein, synteny denotes a minimum of three genes in the vicinity of each other.

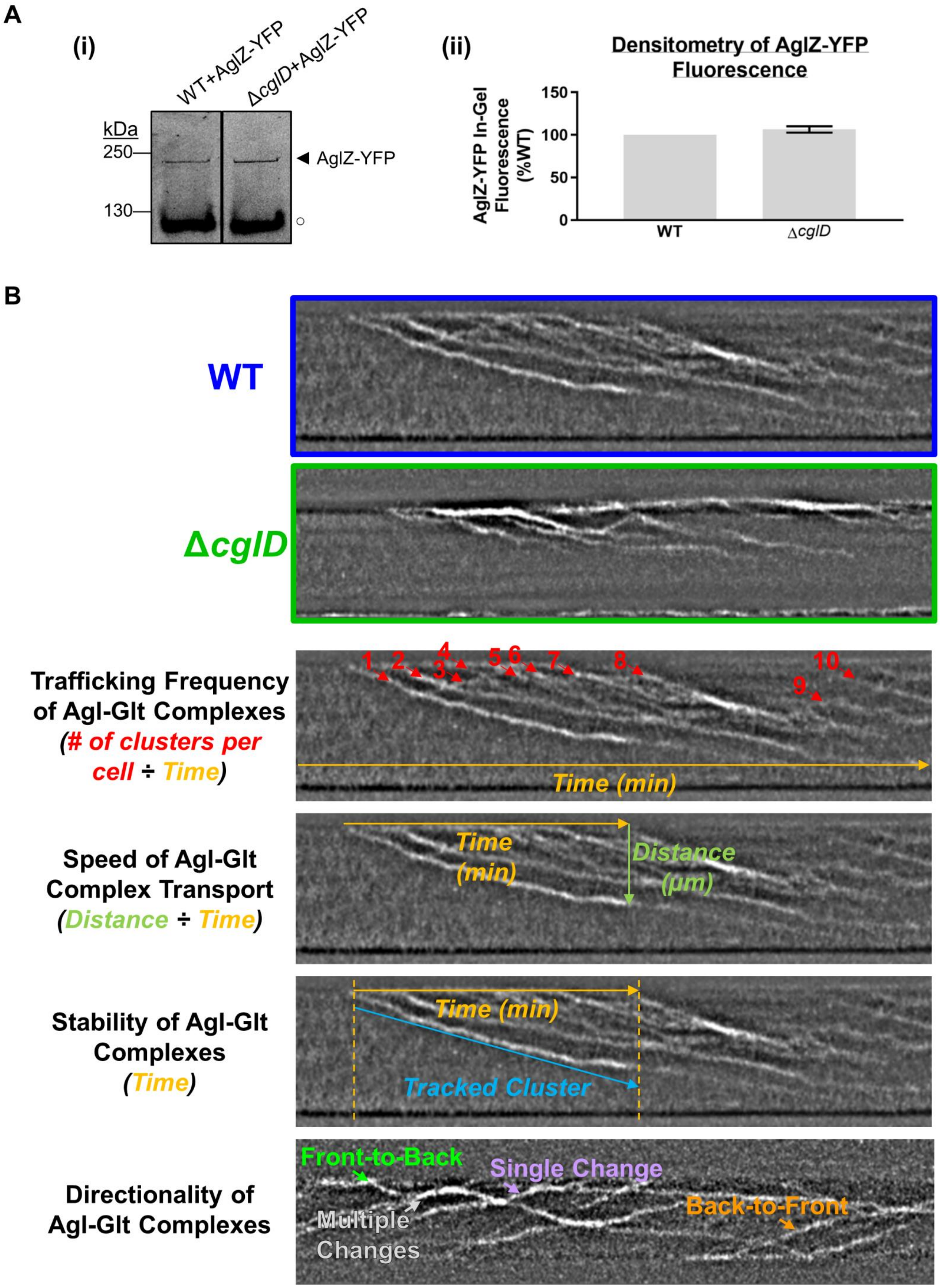
A
(i)
WT+AglZ-YFP
ΔcglD+AglZ-YFP
kDa
250
130
AglZ-YFP
(ii)
Densitometry of AglZ-YFP Fluorescence
AglZ-YFP In-Gel Fluorescence (%WT)
150
100
50
0
WT
ΔcglD
B
WT
ΔcglD
Trafficking Frequency of Agl-Glt Complexes
(# of clusters per cell ÷ Time)
1
2
3
4
5
6
7
8
9
10
Time (min)
Speed of Agl-Glt Complex Transport
(Distance ÷ Time)
Time (min)
Distance (μm)
Stability of Agl-Glt Complexes
(Time)
Time (min)
Tracked Cluster
Directionality of Agl-Glt Complexes
Front-to-Back
Single Change
Multiple Changes
Back-to-Front

**Fig. S6. State of AglZ-YFP in WT vs Δ*cglD* cells.**

**(A)** In-gel fluorescence scan (*panel i*) and densitometry analysis (*panel ii*) of AglZ-YFP levels in WT vs. Δ*cglD* crude-cell lysates resolved via SDS-PAGE. Fluorescence levels were analyzed across five biological replicates and are displayed ± SEM. This difference was not statistically significant, as determined via Wilcoxon signed-rank test performed relative to a reference value of 100 ($p$ value = 0.1875).

**(B)** Kymograph of AglZ-YFP localization in *M. xanthus* cells on chitosan-coated PDMS microfluidic chambers via TIRFM. Arrows in orange denote sequential kymograph slices over time. Arrows in cyan indicate positions of trafficked Agl–Glt clusters in the cell. The manners in which various fluorescent-cluster tracking data (see Fig. 8D-G in the main text) were obtained have been indicated in the example images.

.

**Table S2. Bacterial strains used in this study.**

| **Strain** | **Genotype/Description** | **Construction** | **Source or Reference** |
|---|---|---|---|
| DZ2 | WT | Wild type | 51 |
| TM913 | Δ*cglB* | DZ2 Δ*cglB* | 5 |
| TM490 | Δ*cglD* | DZ2 Δ*cglD* | This work |
| TM600 | Δ*gltK* | DZ2 Δ*gltK* (pBJΔ*gltK*) | 11 |
| TM603 | Δ*gltB* | DZ2 Δ*gltB* (pBJΔ*gltB*) | 11 |
| TM606 | Δ*gltA* | DZ2 Δ*gltA* (pBJΔ*gltA*) | 11 |
| TM570 | Δ*gltC* | DZ2 Δ*gltC* (pBJΔ*gltC*) | 11 |
| TM646 | Δ*gltJ* | DZ2 Δ*gltJ* | 5 |
| TM731 | Δ*gltI* | DZ2 Δ*gltI* | 5 |
| TM149 | Δ*gltH* | DZ2 Δ*gltH* (pBJΔ*gltH*) | 11 |
| TM135 | Δ*gltG* | DZ2 Δ*gltG* (pBJΔ*gltG*) | 11 |
| TM136 | Δ*gltF* | DZ2 Δ*gltF* (pBJΔ*gltF*) | 11 |
| TM148 | Δ*gltE* | DZ2 Δ*gltE* (pBJΔ*gltE*) | 11 |
| TM142 | Δ*gltD* | DZ2 Δ*gltD* (pBJΔ*gltD*) | 11 |
| TM293 | Ω*pilA* | DZ2 Ω*mxan_5783* (Tet$^R$ cassette insertion) | 9 |
| TM913 | Δ*cglB* + Ω*pilA* | DZ2 Δ*cglB* Ω*mxan_5783* (Tet$^R$ cassette insertion) | 5 |
| TM492 | Δ*cglD* + Ω*pilA* | DZ2 Δ*cglD* Ω*mxan_5783* (Tet$^R$ cassette insertion) | This work |
| TM829 | WT + *aglZ-YFP* | DZ2 pBJ*AglZ-YFP* | 6 |
| TM1181 | Δ*cglB* + *aglZ-YFP* | TM913 *aglZ-YFP* (pBJ*AglZ-YFP*) | 5 |
| TM1219 | Δ*cglD* + *aglZ-YFP* | TM490 *aglZ-YFP* (pBJ*AglZ-YFP*) | This work |
| SI101 | WT + *OMss-mCherry* | DZ2 OMss-mCherry (pSWU19-OMss-mCherry) | This work |
| SI102 | Δ*cglB* + *OMss-mCherry* | TM913 OMss-mCherry (pSWU19-OMss-mCherry) | This work |
| SI103 | Δ*cglD* + *OMss-mCherry* | TM490 OMss-mCherry (pSWU19-OMss-mCherry) | This work |
