## Supplementary material for "Integrin-like adhesin CglD confers traction and stabilizes bacterial focal adhesions involved in myxobacterial gliding motility": Table S1

**TABLE S1** **GltD-GST Mass Spectrometry Data**

| Ensembl Gene ID | Gene Symbol | Accessi | Description | Coverage [%] | AvgCoverage | # Peptides | # PSMs | # Unique Peptides | AvgUniPeps | # AAs | MW [kDa] | calc. pI | Score Sequest HT: Sequest HT | AvgScore | # Peptides (by Search Engine): Sequest HT | Biological Process | Cellular Component | Molecular Function | Pfam IDs | KEGG Pathways | # Protein Pathway Groups |
|---|---|---|---|---|---|---|---|---|---|---|---|---|---|---|---|---|---|---|---|---|---|
| **MXAN_0962** | **cglD** | **O1DDC** | **Putative lipoprotein [OS=Myxococcus xanthus DK 1622]** | **33** | **18.5** | **24** | **78** | **24** | **13.5** | **1130** | **117** | **4.37** | **285.55** | **151.835** | **24** | | | | | | **0** |
| MXAN_1623 | MXAN_1623 | O1DBU | peptidase, M16 (pitrilysin) family [OS=Myxococcus xanthus DK 1622] | 29 | 16.5 | 14 | 105 | 14 | 8 | 478 | 53.2 | 5.81 | 327.63 | 173.67 | 14 | | | catalytic activity;metal ion binding | Pf00675, Pf05193 | | 0 |
| MXAN_1624 | MXAN_1624 | O1DBU | peptidase, M16 (pitrilysin) family [OS=Myxococcus xanthus DK 1622] | 27 | 17 | 13 | 194 | 13 | 8 | 473 | 51.1 | 8.5 | 562.25 | 284.22 | 13 | | | catalytic activity;metal ion binding | Pf00675, Pf05193 | | 0 |
| MXAN_1994 | rpsI | O1DAU | 30S ribosomal protein S9 [OS=Myxococcus xanthus DK 1622] | 42 | 31 | 5 | 36 | 5 | 3.5 | 133 | 14.9 | 10.95 | 123.42 | 69.81 | 5 | metabolic process | ribosome | structural molecule activity | Pf00380, Pf02120 | Ribosome | 1 |
| **MXAN_2540** | **gltA** | **O1D9B** | **Uncharacterized protein [OS=Myxococcus xanthus DK 1622]** | **27** | **16** | **4** | **12** | **4** | **2.5** | **256** | **27.8** | **6.35** | **40.8** | **26.975** | **4** | | | | | | **0** |
| MXAN_2674 | MXAN_2674 | O1D8Y | Uncharacterized protein [OS=Myxococcus xanthus DK 1622] | 51 | 35 | 6 | 16 | 6 | 4 | 135 | 14.9 | 10.13 | 48.13 | 30.055 | 6 | | | | | | 0 |
| MXAN_2703 | MXAN_2703 | O1D8V | putative cAMP phosphodiesterases class-II [OS=Myxococcus xanthus DK 1622] | 31 | 20 | 5 | 64 | 5 | 3 | 251 | 27.6 | 7.5 | 214.45 | 108.775 | 5 | metabolic process | | catalytic activity | Pf00753, Pf02112, Pf12706 | | 0 |
| MXAN_2728 | MXAN_2728 | O1D8S | putative NADH dehydrogenase I, G subunit [OS=Myxococcus xanthus DK 1622] | 27 | 22.5 | 10 | 48 | 10 | 8.5 | 541 | 59.3 | 8.68 | 144.93 | 86.73 | 10 | metabolic process | membrane | catalytic activity;metal ion binding | Pf00384, Pf10588, Pf13510 | Oxidative phosphorylation; Metabolic pathways | 2 |
| MXAN_2736 | MXAN_2736 | O1D8S | Uncharacterized protein [OS=Myxococcus xanthus DK 1622] | 41 | 25 | 5 | 262 | 5 | 3 | 78 | 9.1 | 9.17 | 734.79 | 368.16 | 5 | | | | | | 0 |
| MXAN_3160 | MXAN_3160 | O1D7L | peptidase, M13 (neprilysin) family [OS=Myxococcus xanthus DK 1622] | 32 | 17 | 18 | 141 | 18 | 9.5 | 703 | 78.4 | 7.05 | 400.15 | 202.485 | 18 | metabolic process | | catalytic activity | Pf01431, Pf05649 | | 0 |
| MXAN_3581 | MXAN_3581 | O1D6F | Peptidyl-dipeptidase A [OS=Myxococcus xanthus DK 1622] | 34 | 18 | 20 | 80 | 20 | 10.5 | 624 | 70 | 6.8 | 199.2 | 101.725 | 20 | metabolic process | membrane | catalytic activity | Pf01401, Pf10303 | | 0 |
| MXAN_3777 | guaB | O1D5W | inosine-5'-monophosphate dehydrogenase [OS=Myxococcus xanthus DK 1622] | 43 | 23.5 | 17 | 70 | 17 | 9.5 | 485 | 52 | 7.11 | 190.36 | 103.12 | 17 | metabolic process | | catalytic activity;metal ion binding;nucleotide binding | Pf00478, Pf00571, Pf01070, Pf03060 | Metabolic pathways; Purine metabolism; Biosynthesis of secondary metabolites | 3 |
| MXAN_3830 | MXAN_3830 | O1D5R | Uncharacterized protein [OS=Myxococcus xanthus DK 1622] | 24 | 16 | 5 | 23 | 5 | 3 | 325 | 35.9 | 6.23 | 69.78 | 36.2 | 5 | | | | | | 0 |
| MXAN_4635 | MXAN_4635 | O1D3H | Uncharacterized protein [OS=Myxococcus xanthus DK 1622] | 21 | 25 | 5 | 393 | 5 | 5.5 | 240 | 24.7 | 11.12 | 1103.85 | 582.46 | 5 | | | | Pf04519 | | 0 |
| MXAN_4637 | MXAN_4637 | O1D3H | Uncharacterized protein [OS=Myxococcus xanthus DK 1622] | 96 | 66 | 10 | 154 | 10 | 6.5 | 113 | 11.6 | 4.79 | 385.58 | 206.365 | 10 | | | | Pf04519, Pf10988 | | 0 |
| **MXAN_4869** | **gltE (aglT)** | **O1D2U** | **adventurous gliding motility protein AglT [OS=Myxococcus xanthus DK 1622]** | **24** | **16.5** | **9** | **41** | **9** | **6** | **478** | **51.5** | **6.4** | **130.87** | **74.265** | **9** | | | **protein binding** | **Pf00515, Pf13414, Pf13424, Pf13429, Pf13432, Pf14559** | | **0** |
| MXAN_4894 | groES | O1D2S | 10 kDa chaperonin [OS=Myxococcus xanthus DK 1622] | 41 | 25 | 5 | 20 | 5 | 3 | 96 | 10.6 | 7.25 | 54.09 | 27.8 | 5 | metabolic process | cytoplasm | nucleotide binding | Pf00166 | | 0 |
| MXAN_4895 | groEL | O1D2S | 60 kDa chaperonin 2 [OS=Myxococcus xanthus DK 1622] | 70 | 49 | 36 | 404 | 27 | 17.5 | 549 | 57.9 | 5.3 | 1280.21 | 689.33 | 36 | metabolic process | cytoplasm | nucleotide binding | Pf00118 | RNA degradation | 1 |
| MXAN_5040 | MXAN_5040 | O1D2C | Aldehyde dehydrogenase family protein [OS=Myxococcus xanthus DK 1622] | 46 | 24.5 | 20 | 151 | 20 | 11 | 506 | 55.8 | 6.55 | 403.36 | 209.175 | 20 | metabolic process | | catalytic activity | Pf00171 | Biosynthesis of secondary metabolites; Glycolysis / Gluconeogenesis; Pyruvate metabolism; Microbial metabolism in diverse environments; Metabolic pathways | 5 |
| MXAN_5152 | crdB | O1D21 | OmpA family protein [OS=Myxococcus xanthus DK 1622] | 38 | 22 | 6 | 78 | 6 | 4 | 249 | 26.3 | 4.83 | 216.11 | 112.15 | 6 | | membrane | | Pf00691 | | 0 |
| MXAN_5344 | rpsB | O1D1H | 30S ribosomal protein S2 [OS=Myxococcus xanthus DK 1622] | 37 | 20 | 12 | 43 | 12 | 6.5 | 312 | 34.4 | 9.11 | 114.04 | 62.94 | 12 | metabolic process | ribosome | structural molecule activity | Pf00318 | Ribosome | 1 |
| MXAN_5756 | tolB | O1D0D | Protein TolB [OS=Myxococcus xanthus DK 1622] | 23 | 18 | 6 | 14 | 6 | 5 | 431 | 46 | 8.21 | 36.85 | 26.3 | 6 | metabolic process;transport | | catalytic activity;protein binding | Pf00930, Pf04052, Pf07676, Pf10647, Pf14583 | | 0 |
| MXAN_5933 | MXAN_5933 | O1CZV | peptidase, m48 (ste24 endopeptidase) family [OS=Myxococcus xanthus DK 1622] | 36 | 19 | 10 | 53 | 10 | 5.5 | 274 | 29.2 | 9.16 | 140.69 | 71.205 | 10 | metabolic process | | catalytic activity | Pf01435 | | 0 |
| MXAN_6106 | fibA | O1CZD | matrix-associated zinc metalloprotease FibA [OS=Myxococcus xanthus DK 1622] | 28 | 17 | 15 | 131 | 15 | 9.5 | 744 | 79.8 | 5.43 | 424.72 | 226.825 | 15 | metabolic process | | catalytic activity | Pf01447, Pf02868, Pf04151, Pf07504 | | 0 |
| MXAN_6299 | pyk | O1CYU | pyruvate kinase [OS=Myxococcus xanthus DK 1622] | 35 | 18.5 | 13 | 70 | 13 | 7 | 477 | 51.5 | 8 | 180.74 | 95.01 | 13 | metabolic process | | catalytic activity;metal ion binding | Pf00224, Pf02887 | Glycolysis / Gluconeogenesis; Biosynthesis of secondary metabolites; Microbial metabolism in diverse environments; Metabolic pathways; Carbon metabolism; Biosynthesis of amino acids; Pyruvate metabolism; Purine metabolism; Biosynthesis of anti | 9 |
| MXAN_6337 | MXAN_6337 | O1CYR | putative isocitrate dehydrogenase, NAD-dependent [OS=Myxococcus xanthus DK 1622] | 34 | 42.5 | 8 | 183 | 8 | 10.5 | 334 | 35.8 | 6.81 | 579.87 | 523.92 | 8 | metabolic process | | catalytic activity;nucleotide binding | Pf00180, Pf04166 | Biosynthesis of antibiotics; Biosynthesis of secondary metabolites; Citrate cycle (TCA cycle); Biosynthesis of amino acids; Microbial metabolism in diverse environments; Carbon metabolism; Metabolic pathways; 2-Oxocarboxylic acid metabolism | 8 |
| MXAN_6487 | MXAN_6487 | O1CYB | Outer membrane efflux protein domain protein [OS=Myxococcus xanthus DK 1622] | 32 | 19.5 | 13 | 65 | 13 | 8 | 467 | 50.2 | 5.47 | 161.18 | 83.955 | 13 | transport | | transporter activity | Pf02321 | | 0 |
| MXAN_6516 | ahcY | O1CY8 | Adenosylhomocysteinase [OS=Myxococcus xanthus DK 1622] | 46 | 33 | 29 | 693 | 29 | 18.5 | 476 | 52.3 | 6.54 | 2043.52 | 1093.335 | 29 | metabolic process;transport | cytoplasm | catalytic activity;nucleotide binding | Pf00670, Pf01262, Pf02254, Pf02826, Pf05221 | Metabolic pathways; Cysteine and methionine metabolism | 2 |
| MXAN_6660 | MXAN_6660 | O1CXU | Uncharacterized protein [OS=Myxococcus xanthus DK 1622] | 34 | 26.5 | 8 | 36 | 8 | 5.5 | 250 | 25.3 | 4.6 | 93.9 | 57.14 | 8 | | | | | | 0 |
| MXAN_6720 | MXAN_6720 | O1CXN | Putative lipoprotein [OS=Myxococcus xanthus DK 1622] | 21 | 18 | 3 | 693 | 3 | 2.5 | 115 | 11.4 | 6.52 | 2437.07 | 1294.49 | 3 | | | | | | 0 |
| MXAN_6829 | MXAN_6829 | O1CXC | putative late embryogenesis abundant-like protein [OS=Myxococcus xanthus DK 1622] | 29 | 16.5 | 5 | 28 | 5 | 3 | 256 | 26.9 | 9.85 | 79.21 | 41.98 | 5 | response to stimulus | | | Pf03168 | | 0 |
| MXAN_7212 | MXAN_7212 | O1CW9 | Uncharacterized protein [OS=Myxococcus xanthus DK 1622] | 56 | 42.5 | 7 | 30 | 7 | 5 | 148 | 14.8 | 6.54 | 93.35 | 58.945 | 7 | | | | | | 0 |
